# Tissue-intrinsic Wnt signals antagonize Nodal-driven AVE differentiation

**DOI:** 10.1101/2023.05.19.541432

**Authors:** Sina Schumacher, Max Fernkorn, Michelle Marten, Yung Su Kim, Ivan Bedzhov, Christian Schröter

## Abstract

The anterior-posterior axis of the mammalian embryo is laid down by the anterior visceral endoderm (AVE), an extraembryonic signaling center that is specified within the visceral endoderm. Current models posit that AVE differentiation is promoted globally by epiblast-derived Nodal signals, and spatially restricted by a BMP gradient established by the extraembryonic ectoderm. Here, we report spatially restricted AVE differentiation in bilayered embryo-like aggregates made from mouse embryonic stem cells that lack an extraembryonic ectoderm. Notably, clusters of AVE cells also form in pure visceral endoderm cultures upon activation of Nodal signaling, indicating that tissue-intrinsic factors restrict AVE differentiation. We identify Wnt signaling as a tissue-intrinsic factor that antagonizes AVE-inducing Nodal signals. Together, our results suggest that interactions between epiblast and visceral endoderm alone enable local AVE differentiation in the absence of graded BMP signals. This may be a flexible solution for axis patterning in a wide range of embryo geometries.

## Introduction

Identifying cell-cell communication mechanisms that orchestrate the self-organized development of the mammalian embryo is a major goal in developmental biology. The modularity of stem cell-based embryo-like models offers the possibility to investigate cell differentiation in subsystems, and thereby to identify signaling mechanisms that may have remained hidden in the embryo.

One of the most fundamental processes in embryonic development is the establishment of an anterior-posterior axis. In mammals, this axis is laid down by the anterior visceral endoderm (AVE), a specialized extraembryonic cell population within the visceral endoderm (VE) that overlies the embryonic epiblast at the time of implantation. The AVE expresses transcription factors such as Otx2, Eomes, Gsc and Lhx1, and Wnt, BMP and Nodal antagonists such as Dkk1, Cer1, and Lefty1.^1–3^ These secreted signaling antagonists pattern the epiblast by restricting Wnt, BMP and Nodal signaling to its posterior end, thereby establishing the anterior-posterior axis of the embryo.

In rodents, the VE and the epiblast form a cup-shaped egg cylinder (Figure 1A). The precursor cells of the AVE initially differentiate from the VE at the distal tip of the egg cylinder, before migrating towards the future anterior side. AVE differentiation is promoted by Nodal signals from the epiblast,^4^ and thought to be locally restricted by graded inhibitory BMP signals from the extraembryonic ectoderm (ExE), an extraembryonic tissue at the proximal end of the egg cylinder.^3, 5, 6^ Cell populations with a similarity to the mouse AVE have been described in non-rodent mammals, including humans.^7, 8^ Embryos from these species are disc-rather than cup-shaped, and may therefore lack the BMP gradient present in rodent embryos. This raises the possibility that alternative mechanisms for AVE differentiation exist that may be obscured by the activity of graded BMP signals in rodent embryos.

**Figure 1.**
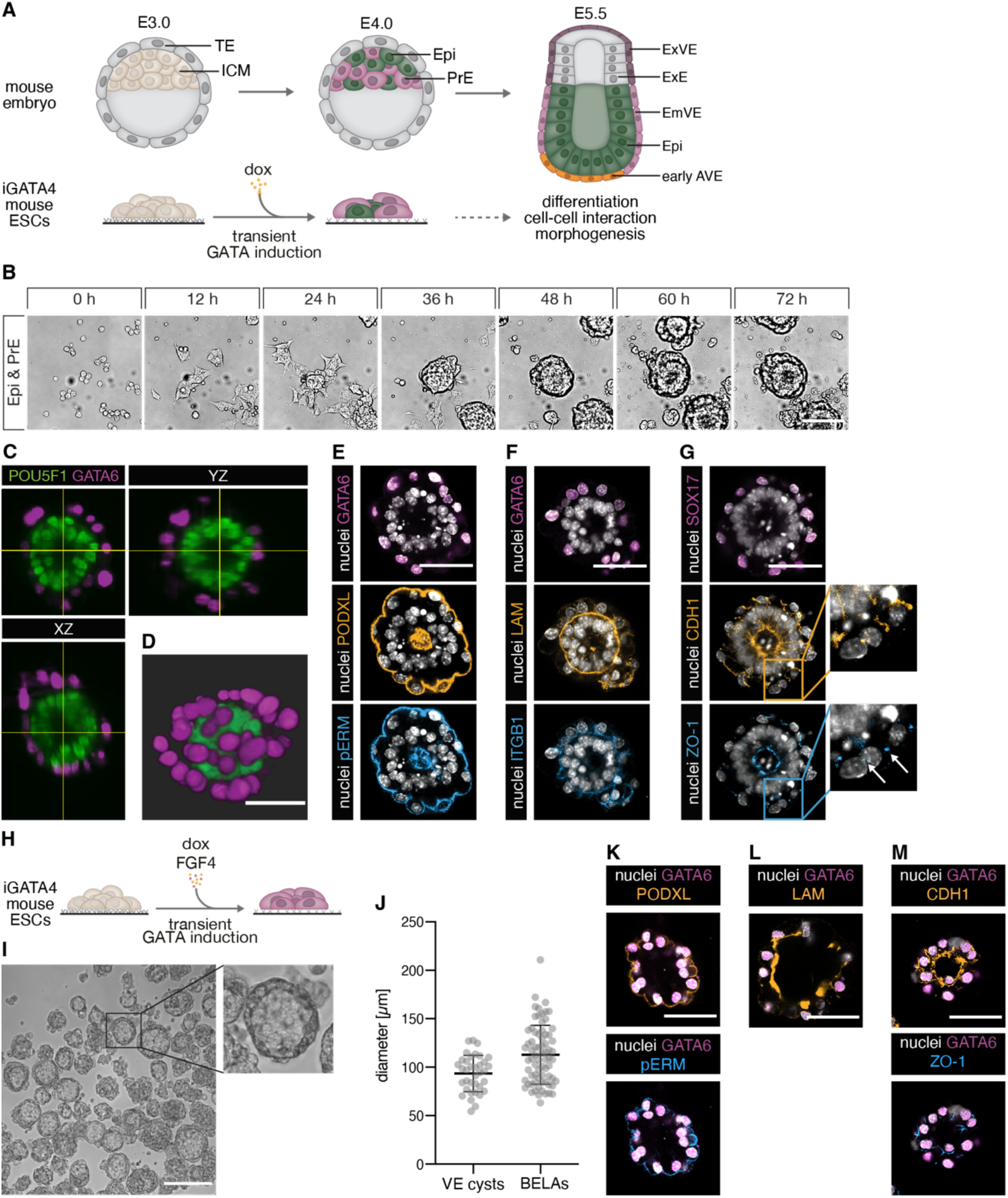
Formation and characterization of BELAs and VE cysts. (A) Schematic of mouse embryonic development from E3.0 to E5.5 (top) and GATA4-inducible embryonic stem cell (ESC) system to model interactions between epiblast (Epi) and extraembryonic endoderm (bottom). (B) Stills from a movie of ESC-derived Epi and primitive endoderm (PrE) cells seeded on a low adhesion substrate in N2B27 medium. (C and D) Orthogonal views (C) and 3D volume rendering (D) of a bilayered aggregate imaged with light sheet microscopy. POU5F1 (green) marks Epi identity and GATA6 (magenta) marks PrE/VE identity. See also Video S2. (E-G) Immunostainings of bilayered aggregates for the PrE/VE markers GATA6 (E, F) or SOX17 (magenta, G), the apical markers PODXL (orange) and pERM (blue) (E), the basement membrane and adhesion markers LAM (orange) and ITGB1 (blue) (F), and the epithelial markers CDH1 (orange) and ZO-1 (blue) (G). Arrows in (G, inset) mark punctate ZO-1 staining characteristic for tight junctions. (H) Schematic of experimental protocol to differentiate pure populations of PrE cells. (I) VE cysts formed in N2B27 supplemented with FGF4 on a low adhesive substrate. (J) Diameters of detached BELAs and VE cysts grown for three days on a low adhesive substrate. n = 72 (BELAs) and n = 36 (VE cysts); bars indicate mean ± SD. (K-M) Immunostainings of VE cysts for the same markers as in (E-G). Scale bars: 50 µm in (B, D-G, and K-M), 200 μm in (H).

Such alternative mechanisms can be identified with stem cell-based embryo models composed of embryonic and specific extraembryonic lineages. Here, we use an embryo-like model system consisting of the epiblast and the VE compartment to study how cell-cell communication controls AVE differentiation in the absence of an ExE. We first characterize bilayered aggregates made from mouse embryonic stem cells (ESCs) that recapitulate the interaction of the epiblast and the VE lineage as seen in the embryo, and contrast them with 3D structures that consist of either of the two cell types alone. Using single-cell RNA-sequencing, we show that the presence of the epiblast compartment suffices to trigger differentiation of a subset of VE cells towards an AVE identity. We apply cell-cell communication analysis to identify the associated signaling pathways, and use this knowledge to develop protocols for AVE differentiation in the absence of an epiblast compartment. Stimulation of Activin/Nodal signaling, coupled to inhibition of Wnt signaling allows us to differentiate almost pure populations of AVE cells in vitro, suggesting that tissue-intrinsic Wnt signals restrict AVE differentiation to local cell clusters. These new signaling mechanisms for AVE differentiation may help explain axis patterning in embryos that do not have a BMP gradient.

## Results

### Generation of simplified 3D models of the Epi- and VE-compartments

To generate a 3D model of the peri-implantation embryo that contains its epiblast and VE-compartment we started from GATA4-inducible mouse ESCs. We have previously shown that following transient GATA4 expression, these cells differentiate into robust proportions of epiblast (Epi) and primitive endoderm (PrE) cells, the precursors of the VE (Figure 1A).^9, 10^ To promote cell-cell interactions, we re-seeded these cell type mixtures after 16 h and lowered the adhesiveness of the substrate. Under these conditions, cells quickly aggregated, formed round structures consisting of two layers of cells that surrounded a central lumen, and eventually detached from the culture surface (Figure 1B; Video S1). The outer layer of these spherical structures consisted of GATA6-positive VE cells, while the inner layer expressed the Epi marker POU5F1 (Figures 1C and 1D; Video S2). Staining with the apical markers PODXL and pERM showed that both compartments were polarized, with the apical domain of the VE pointing towards the outside of the aggregates, and the apical domain of the Epi layer pointing to the inside (Figure 1E). At their basal sides, we detected expression of β1-integrin (ITGB1) as well as a laminin-rich basal membrane (Figure 1F; Video S3). Both layers stained positive for the epithelial markers E-Cadherin (CDH1) and ZO-1 (Figure 1G). This architecture of two apposed epithelial layers resembles the arrangement of cells in the distal part of the egg cylinder. We therefore term these structures “bilayered embryo-like aggregates” (BELAs).

We next sought to generate 3D structures that consist of each of the two cell types in isolation. To obtain only Epi cells, we cultured ESCs under the same conditions as used for BELA formation, but omitted the doxycycline pulse. Under these conditions we observed extensive cell death from 48 h after re-seeding onwards (Figures S1A-S1C). As previously described, culture of cells in matrigel rescued their survival and induced cyst formation (Figure S1D),^11^ suggesting that a major function of the VE layer in BELAs is to provide survival and patterning signals via the extracellular matrix.

To generate pure cultures of PrE cells, we extended the expression of the inducible transgene for 16 h after the switch to N2B27 medium, and supplemented the medium with exogenous FGF4 (Figures 1H and S1A).^9^ When these cultures were re-seeded on low-adhesion substrates in N2B27 only, we observed rapid cell death (Figures S1B and S1C), but survival could be fully rescued by continued addition of FGF4 (Figure S1B). Surprisingly, in the presence of FGF4, these cells aggregated and formed non-adherent 3D structures (Figures 1I and S1B). These aggregates varied in size and shape, but a large number of them formed round cysts with a big lumen (Figure 1I). The diameter of these cysts was 93.4±18.8 µm (mean ± SD), similar to that of BELAs (112.8±30.4 µm, Figure 1J). The apical markers PODXL and pERM localized to the outside of the cysts, laminin was secreted to their inside, and the localized expression of CDH1 and ZO-1 further indicated an epithelial organization (Figures 1K-1M). Thus, these structures resemble the outer layer of BELAs, and we hence refer to them as VE cysts.

Taken together, the exchange of mutual survival signals between Epi and PrE cells underlies the spontaneous formation of BELAs. Replacing these signals with purified factors allows us to generate Epi- and VE cysts that consist of only one of the cell types found in BELAs, but that capture the 3D organization of the single compartments.

### Interactions between Epi and VE cells in BELAs shape cell differentiation trajectories

In the post-implantation embryo, differentiation of both the VE as well as the Epi lineage is strongly influenced by signals from the ExE. We reasoned that the three simplified 3D models could reveal mechanisms for cell differentiation that are independent from the ExE. We therefore performed single-cell RNA-sequencing (scRNAseq) on the three types of aggregates (Figure 2A). Representation of the single-cell transcriptomes in a UMAP plot showed two major groups, one containing mainly cells from Epi cysts and a subgroup of the BELA cells. The other group contained the remaining BELA cells as well as most cells from VE cysts (Figure 2B). Expression of the VE marker genes *Sox17*, *Cubn, Dab2*, and *Gata6*^11^ and the Epi marker genes *Fgf4, Nanog*, *Pou5f1*, and *Sox2*^12^ identified these two broad groups as VE and Epi, respectively (Figure 2C). To determine which cell types and developmental stages were captured in the in vitro samples, we integrated our scRNAseq data with single-cell transcriptomes from the embryo. We chose a reference dataset that covered several embryonic stages between E3.5 and E8.75, and that focused in particular on the emergence of the endoderm lineage.^13^ UMAP representations after integration indicated that cells from BELAs and cysts corresponded to a range of embryo cell types (Figures 2D and 2E).

**Figure 2.**
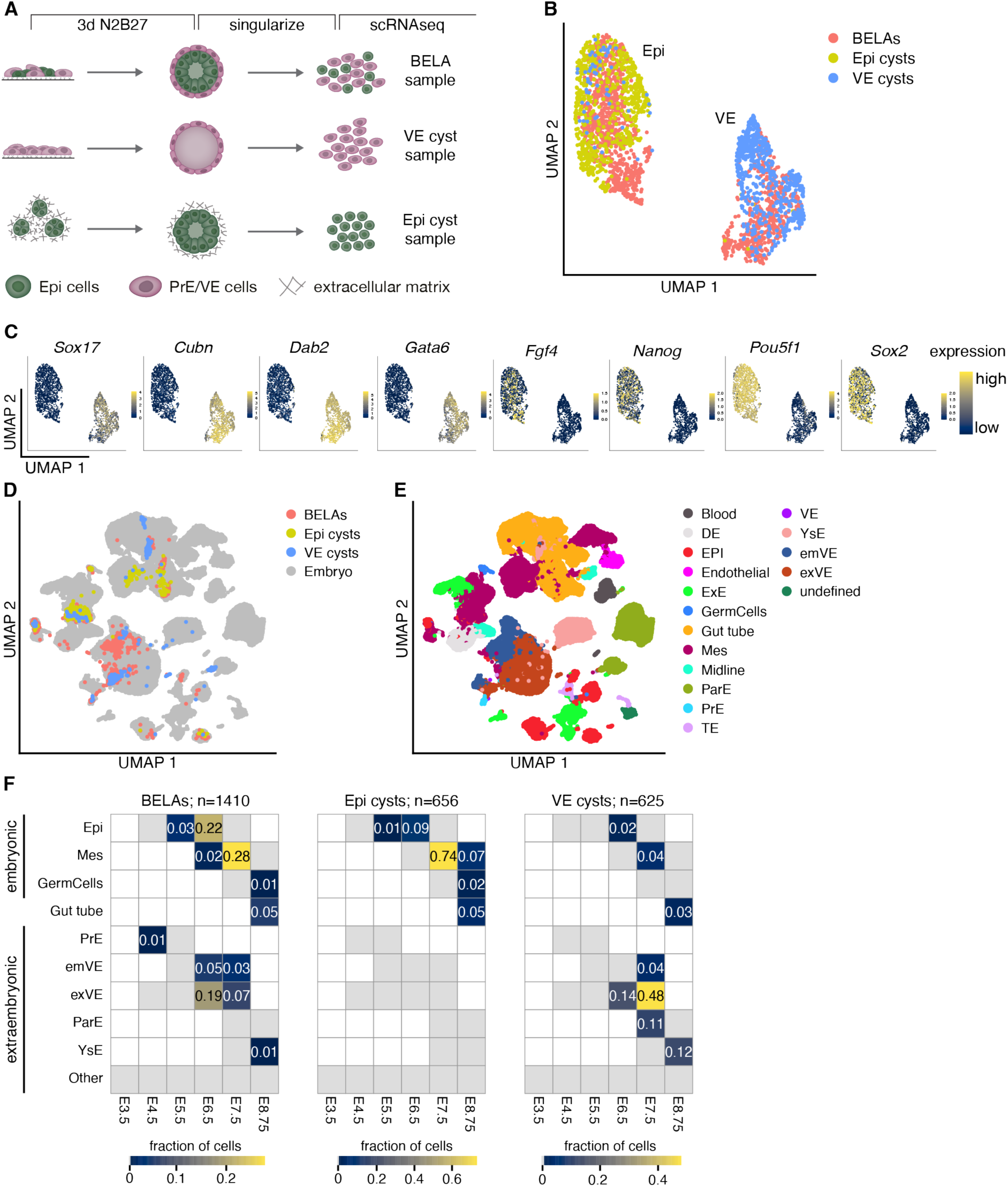
Single-cell RNA-sequencing and data integration to determine cell type identities in BELAs and cysts. (A) Experimental approach to prepare samples for scRNAseq. (B) UMAP of batch corrected single-cell transcriptomes from cells prepared as in (A). Colors indicate sample of origin. (C) Expression levels of VE markers Gata6, Sox17, Dab2, and Cubn, and Epi markers Pou5f1, Sox2, Nanog, and Fgf4. To better visualize the cell type-specific expression of Fgf4, Nanog and Sox2, expression levels above log2 ≥ 1.5 (Fgf4) or log2 ≥ 2 (Nanog and Sox2) are shown in yellow. (D) UMAP of single-cell transcriptomes from BELAs, Epi cysts and VE cysts, integrated with scRNAseq data from mouse embryos covering stages E4.5 to E8.75 (Nowotschin et al., 2019).^13^ (E) Same UMAP as in (D), colored according to cell type annotation from Nowotschin et al., 2019 after integration and label transfer. (F) Heatmaps showing the fraction of cells in BELAs (left), Epi cysts (middle) and VE cysts (right) assigned to particular cell types and time points from the embryo. Because the E8.75 gut tube has both embryonic and extraembryonic origin,^14, 15^ it was not assigned to any of the two categories.

We transferred cell type and stage labels from the reference dataset and plotted their frequency in each sample (Figure 2F). While cells from BELAs mapped to both E6.5 and E7.5 reference cells, the majority of cells from both Epi and VE cysts mapped to E7.5, indicating that cyst cells were developmentally more advanced than cells from BELAs. BELA cells mapped to both embryonic cell types (Epi, mesoderm (Mes), and germ cells) and extraembryonic cell types (PrE, embryonic VE (emVE), extraembryonic VE (exVE), parietal endoderm (ParE), and yolk sac endoderm (YsE)). Cells from Epi cysts in contrast mapped mostly to embryonic cell types, whereas cells from VE cysts mapped to extraembryonic cell types. The low number of cells from VE cysts that mapped to embryonic cell types likely originate from a small fraction of cells is refractory to PrE differentiation because of insufficient transgene induction levels.^9^ The vast majority of embryonic cells from Epi cysts were labeled as mesoderm, whereas the embryonic cells from BELAs were labeled both as Epi and mesoderm. The extraembryonic cells from VE cysts mostly mapped to cell types that are not in contact with the epiblast, such as the exVE, the ParE, and the YsE, and only 4% mapped to the emVE. In BELAs in contrast, 8% of all cells were labeled as emVE, corresponding to approximately one fifth of all extraembryonic cells in this sample, and indicating that the presence of the Epi core in BELAs promotes an emVE identity. We conclude that cells from all three 3D systems bear transcriptional similarity to the embryonic and extraembryonic lineages of the mouse embryo shortly after implantation. Furthermore, differences in developmental stage and cell type identity between embryonic cells from Epi cysts and BELAs, and between extraembryonic cells from VE cysts and BELAs, indicate that interactions between the two cell types regulate cell differentiation.

### Interaction of Epi and VE cells in BELAs promotes AVE differentiation

To characterize in more detail how interactions between Epi and VE cells in BELAs affect cell differentiation, we clustered the single-cell transcriptomes and searched for cell types that were present in BELAs, but not in the cyst samples. Epi cells clustered according to their sample of origin, with one cluster containing the Epi cells from BELAs (cluster 1), and the other one containing almost all Epi cyst cells (cluster 2, Figures 3A and 3B). These global transcriptomic differences between Epi cells from the two sample types suggests that the signaling environment generated by the VE layer in BELAs differs from that generated by the artificial extracellular matrix used to grow Epi cysts. VE cells also fell into two clusters, but here, cells were not segregated based on their origin (Figures 3A and 3B). Instead, cluster 3 consisted of both cells from VE cysts and BELAs, whereas a small cluster 4 contained exclusively cells from BELAs (Figures 3A and 3B). Genes that were downregulated in cluster 4 mostly encoded components of the extracellular matrix (Figure S2, Table S1). The list of upregulated genes on the other hand contained markers such as Lhx1, Otx2, Eomes and Gsc, suggesting that cells in this cluster had adopted an AVE identity (Figure 3C, Table S1). To corroborate this finding, we integrated the transcriptomes of BELA cells from clusters 3 and 4 with an embryo data set that focused especially on the AVE differentiation from E5.5 to 6.25.^16^ Consistent with the integration with the whole-embryo dataset above, the majority of cells from both cluster 3 and 4 mapped together with E6.25 embryo cells (79% and 76%, respectively, Figures 3D-3F). While 95% of cells from cluster 4 were labeled as AVE after integration, cells from cluster 3 were labeled both as AVE (39%) and Epi-VE (59%, Figure 3F). This supports the notion of AVE differentiation in BELAs, and suggests that AVE gene expression signatures extend to cells beyond those identified in cluster 4. In contrast to integration with the whole-embryo dataset, where a large proportion of BELA-VE cells were mapped to the exVE lineage, virtually no cells obtained the corresponding ExE-VE label upon integration with the AVE-focused dataset. These discrepancies could be due to diverging strategies for annotation in the two reference datasets, as well as different representations of the lineages, which may bias the outcome of the dataset integration.

**Figure 3.**
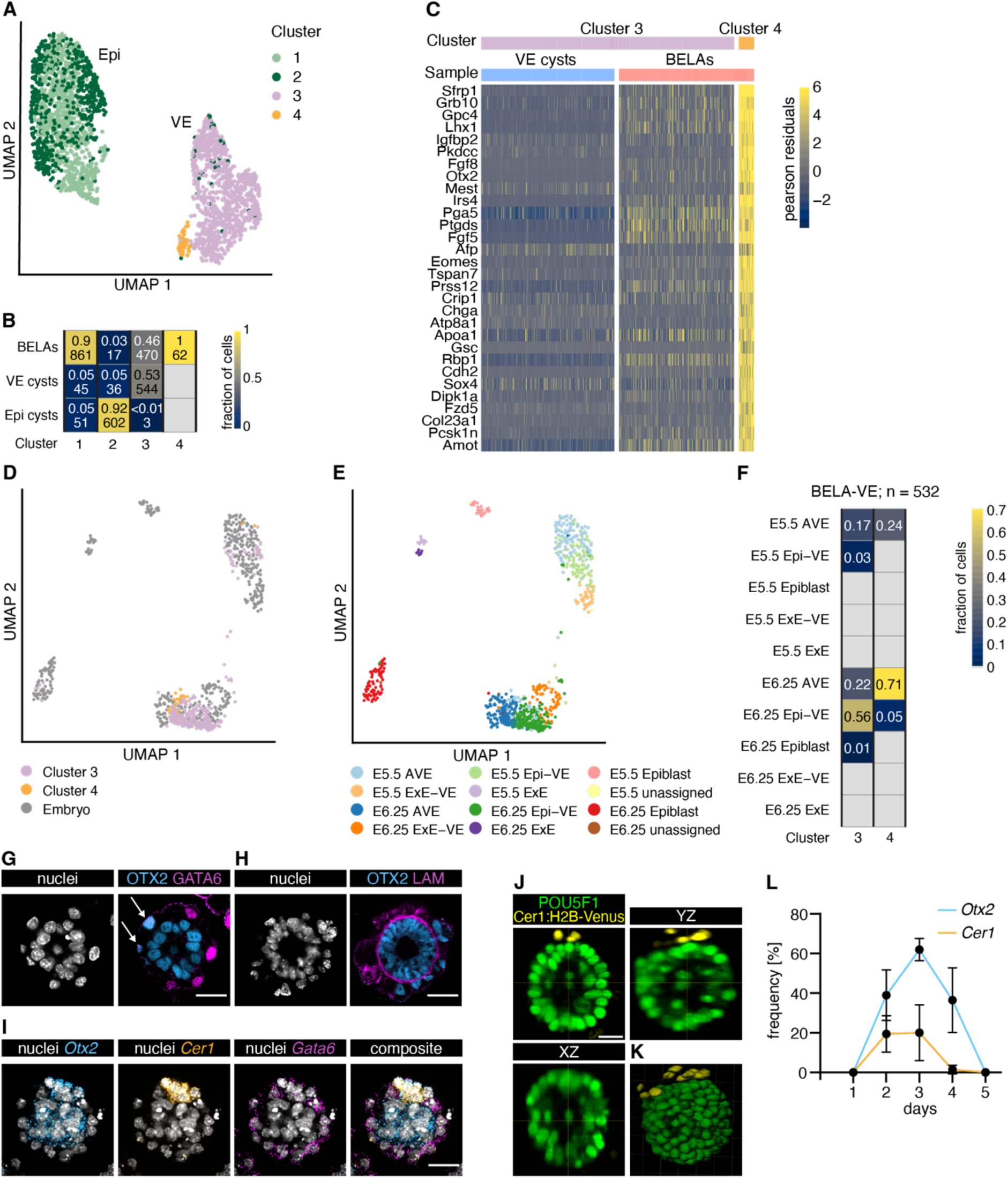
AVE differentiation in BELAs. (A) UMAP representation of single-cell transcriptomes (same as in Figure 2B), colored according to Louvain clustering. (B) Heatmap showing the fraction and total number of cells from each sample in the four clusters from (A). The small number of cells from VE cysts in clusters 1 and 2 likely originate from cells that were refractory to PrE differentiation (see above, and Raina et al., 2021^9^). (C) Heatmap showing the 30 most upregulated genes between the cells of cluster 3 and cluster 4 in (A), ordered by log2-fold change. Single-cell expression is shown as the Pearson residual of the normalized counts. (D) UMAP of single-cell transcriptomes from BELA-VE cells (Cluster 3 and Cluster 4 in (A), (B)), integrated with scRNAseq data from mouse embryos at E5.5 and E6.25 from Thowfeequ et al., 2021^16^. (E) Same UMAP as in (D), colored according to cell type annotation from Thowfeequ et al., 2021^16^ after integration and label transfer. (F) Heatmap showing the fraction of BELA-VE cells assigned to particular cell types and developmental time points from the embryo. (G) Immunostaining for the AVE marker OTX2 (blue) and the VE marker GATA6 (magenta). Arrows highlight co-expression. (H) Immunostaining for the AVE marker OTX2 (blue) and the basement membrane marker LAM (magenta). (I) In situ HCR staining for the AVE markers *Otx2* (blue) and *Cer1* (orange), and the VE marker *Gata6* (magenta). (J and K) Orthogonal views (J) and 3D volume rendering (K) of a BELA stained for the Epi marker POU5F1 (green) and the AVE reporter Cer1:H2B-Venus (yellow) imaged with light sheet microscopy. (L) Mean frequency of AVE marker gene expression in BELAs on different days after re-seeding. *Otx2* expression was scored as AVE marker only if it could clearly be assigned to the outer layer of BELAs. N = 2, n ≥ 18, except for day 1 n = 10, error bars indicate SD. Scale bars: 25 µm.

To validate the presence of AVE cells, and to determine how they are distributed amongst individual BELAs, we visualized expression of the AVE markers *Otx2* and *Cer1* together with the pan-VE marker *Gata6*. In 28 out of 33 BELAs, we found cells that co-expressed OTX2 and GATA6 protein (Figure 3G). Furthermore, such OTX2-positive cells were located outside the laminin-ring, as would be expected for an AVE identity (Figure 3H). Similar results were obtained by in situ HCR staining for *Otx2* and *Cer1* mRNA (Figure 3I, 27 out of 36 BELAs with cells co-expressing *Otx2* and *Gata6* mRNA; 10 out of 36 BELAs with cells co-expressing *Cer1* and *Gata6* mRNA). Light-sheet imaging of a Cer1:H2B-Venus transcriptional reporter^17^ (Figure S3) integrated into our inducible lines indicated that AVE cells in BELAs tended to be spatially clustered (Figure 3J and 3K, Video S4). In the embryo, AVE markers such as Otx2 and Cer1 cells are only transiently expressed between E5.5 and E7.5.^13^ BELAs expressing these markers in the VE could first be detected two days after re-seeding, their number peaked at day 3, and declined thereafter (Figure 3L), thus recapitulating the transient nature of the AVE in the embryo. Taken together, these results show that interactions between the Epi and the VE trigger AVE differentiation in small groups of spatially clustered cells in a large number of BELAs.

### Activin/Nodal signaling is necessary and sufficient for AVE differentiation

Next, we used LIANA,^18^ a ligand-receptor analysis framework, to identify potential Epi-derived signals that could trigger AVE differentiation in BELAs. Amongst the top scoring interactions between Epi and VE cells, we found ligand-receptor pairs associated with signaling from the extracellular matrix, FGF, and Eph-Ephrin signaling (Figure 4A, Table S2). Consistent with the critical role of Nodal signaling for AVE differentiation in the embryo,^4^ this analysis furthermore returned the Nodal receptors Acvr1b and Acvr2a and Nodal co-factor ligand Tdgf1. To test the function of Nodal signaling in BELAs, we used the receptor inhibitor SB431542 (SB43), and analyzed AVE differentiation in BELAs generated from Nodal mutant cells. Both perturbations abrogated AVE differentiation, as judged by Otx2 and Cer1 expression (Figure 4B-4D). Epiblast-derived Nodal signals were required for AVE differentiation not only in BELAs but also in the embryo, since tetraploid complementation with Nodal mutant cells likewise resulted in the absence of a CER1-positive AVE (Figure 4E).

**Figure 4.**
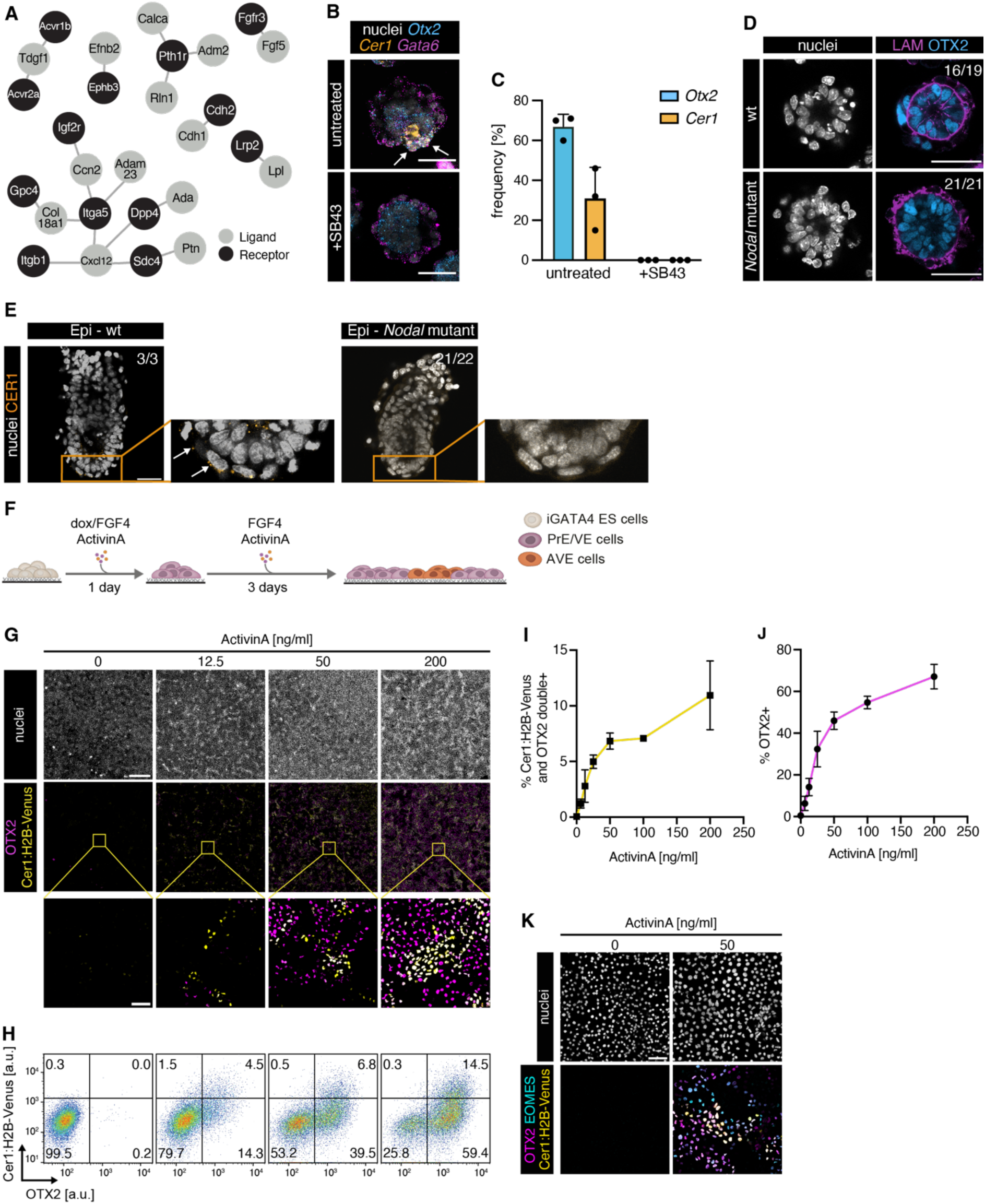
Activin/Nodal signaling is necessary and sufficient for AVE differentiation. (A) Output of ligand-receptor analysis with LIANA,^18^ showing the top 20 interactions between Epi-derived ligands and VE-derived receptors. (B) In situ HCR staining of untreated (top) and SB43-treated (bottom) BELAs for AVE markers *Otx2* (blue) and *Cer1* (orange) and the PrE/VE marker *Gata6* (magenta). (C) Mean frequency of AVE marker gene expression in untreated and SB43-treated BELAs three days after re-seeding. N = 3, n ≥ 20 per condition, error bars indicate SD. (D) Immunostaining for Laminin (magenta) and OTX2 (blue) in BELAs made from *Nodal* wild-type (top) and *Nodal*-mutant cells (bottom). (E) Immunostaining for CER1 (orange) in E5.5 mouse embryos generated via tetraploid complementation with wild-type (left) or *Nodal-*mutant cells (right). (F) Schematic of experimental protocol to generate 2D layers of VE cells for AVE differentiation. (G) Immunostaining for OTX2 (magenta) and H2B-Venus (yellow) of Cer1:H2B-Venus reporter cells treated with indicated concentrations of ActivinA for 3 days after an extended doxycycline pulse. (H) Flow cytometry of cells differentiated and stained as in (G). (I) Mean percentage of Cer1:H2B-Venus; OTX2 double-positive cells differentiated with increasing doses of ActivinA. N = 3, error bars indicate SD. (J) Same as (I) but showing percentage of OTX2-positive cells. (K) Immunostaining for OTX2 (magenta), EOMES (cyan), and H2B-Venus (yellow) of Cer1:H2B-Venus reporter cells treated as in (G). Scale bars: 50 µm in (B), (D), (E), ((G) inset), (K); 500 µm in (G).

We then asked whether Epi-derived Nodal signals were sufficient to trigger AVE differentiation in vitro. We again used an extended doxycycline pulse together with exogenous FGF4 to generate pure cultures of PrE cells, and seeded these on a high-adhesion substrate to analyze differentiation in a homogeneous 2D VE-layer (Figure 4F). Addition of the Nodal agonist ActivinA triggered the expression of both the Cer1:H2B-Venus reporter, as well as OTX2 expression, in a dose-dependent manner (Figures 4G-4J). Cultures were homogeneously GATA6-positive, both in the presence and absence of ActivinA (Figure S4). At 200 ng/ml ActivinA, approximately two thirds of all cells expressed OTX2, indicating that the majority of VE cells have AVE differentiation potential. ActivinA also triggered expression of the AVE marker EOMES (Figure 4K). Intriguingly, cells expressing AVE markers often occurred in spatial clusters, with a central Cer1:H2B-Venus/OTX2/EOMES triple-positive core, surrounded by cells that were OTX2 and EOMES single- or double-positive (Figure 4G and 4K). Taken together, these experiments show that Activin/Nodal signals are necessary and sufficient for the differentiation of AVE cells.

### Wnt signaling restricts AVE differentiation to local cell clusters

We then wondered why AVE differentiation occurred in spatial clusters despite global stimulation with ActivinA in homogeneous 2D layers of VE cells. This could reflect clonal expansion of single cells that were privileged for AVE differentiation, or alternatively, be the consequence of local signaling domains that allow for AVE differentiation. To distinguish between these possibilities, we added three different fluorescent labels to the inducible cell lines and analyzed the clonal composition of Cer1:H2B-positive nests in mixed cultures (Figures 5A and 5B). We found a similar number of nests that carried the same clonal label (13/30) and nests composed of cells with different labels (17/30, Figure 5B). This suggests that the clonal expansion of single cells contributes to nest formation, but that in addition, local signaling environments generated by cell-cell communication promote AVE differentiation. In the embryo, BMP4 signals restrict differentiation.^6^ However, addition of BMP4 only mildly reduced the proportion of Cer1:H2B-Venus-positive cells and had no effect on OTX2 expression (Figure S5). Addition of the BMP receptor inhibitor LDN193189 did not increase the expression of either of the two markers (Figure S5). Therefore, BMP signaling does not play a strong role in restricting AVE differentiation in vitro. Since we noticed that Nodal was specifically expressed in AVE cells in BELAs (Figure S6A), we reasoned that a positive feedback loop centered on Nodal signaling could promote AVE differentiation in nests. When we measured expression of AVE markers in Nodal-mutant cells, we found that the proportion of Cer1:H2B-Venus-positive cells was reduced by half compared to wild-type controls, but the proportion of OTX2-positive cells was unchanged (Figures S6B-S6F), and cells expressing these AVE markers were still spatially clustered (Figure S6D). Therefore, endogenous Nodal signaling plays a minor role in regulating AVE differentiation in vitro.

**Figure 5.**
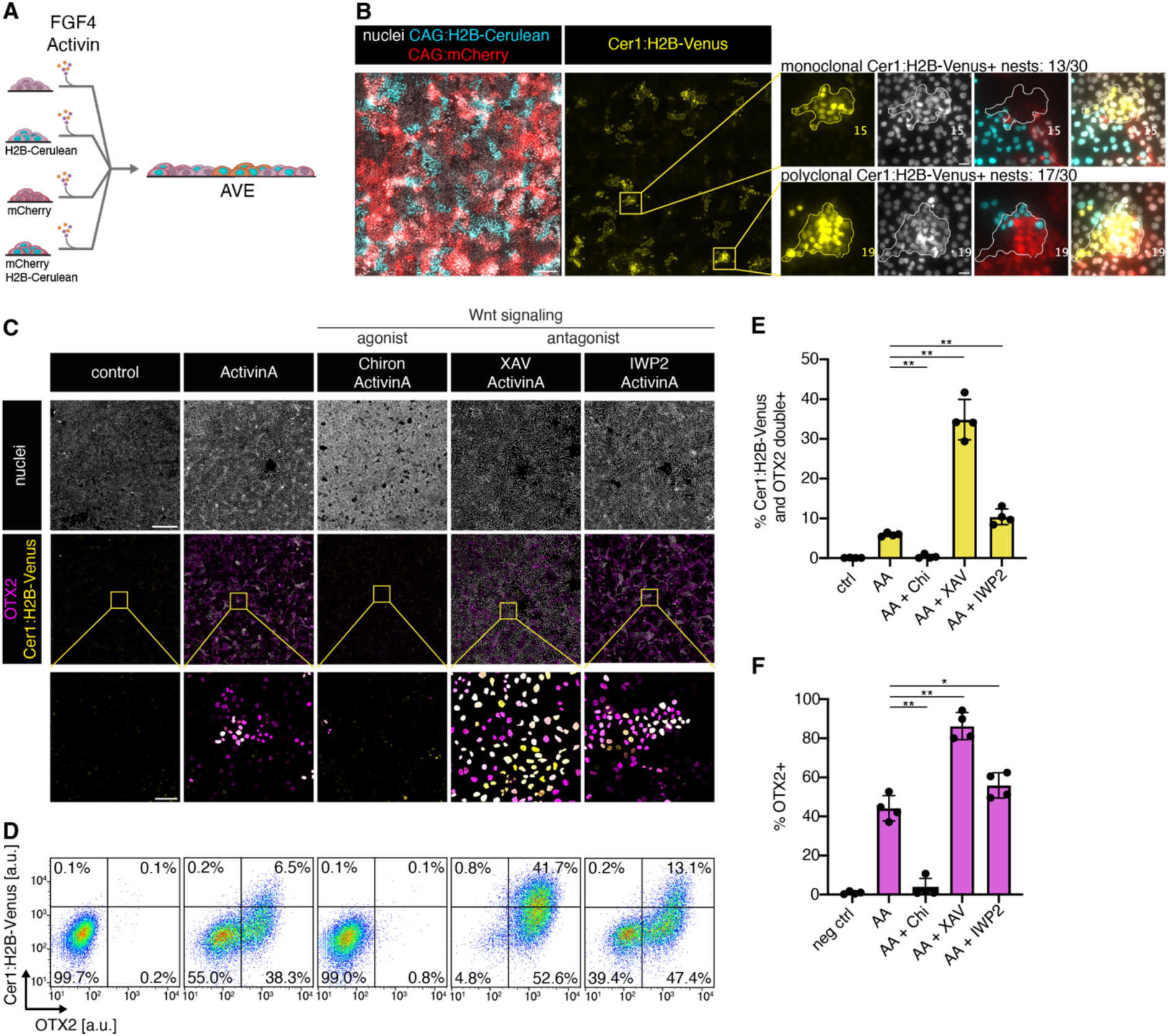
Tissue-intrinsic Wnt signaling regulates AVE differentiation. (A) Experimental approach to determine clonal composition of AVE nests. (B) Expression of clonal labels (red, cyan) and Cer1:H2B-Venus (yellow) reporter in cultures differentiated as in (A). Insets on the right show examples of Cer1:H2B-Venus-expressing nests with a single clonal label (top, 13/30 nests), or with multiple labels (bottom, 17/30 nests). (C) Immunostaining for OTX2 (magenta) and H2B-Venus (yellow) of Cer1:H2B-Venus reporter cells differentiated for 3 days after an extended doxycycline pulse with 50 ng/ml ActivinA (AA), together with 3 µM Chir99021 (Chiron), 20 µM XAV939 (XAV), or 2 µM IWP2 as indicated. (D) Flow cytometry of cells differentiated and stained as in (C). (E) Mean percentage of Cer1:H2B-Venus; OTX2 double-positive cells differentiated as in (C). N = 4, error bars indicate SD. (F) Same as (E) but showing percentage of OTX2-positive cells. Scale bars: 200 µm ((B) overview); 20 µm ((B) inset); 500 µm ((C) overview); 50 µm ((C) inset). * and ** in (E), (F) indicate p ≤ 0.05 and p ≤ 0.005 determined by a two-tailed, unpaired t-test, respectively.

Finally, given the important role of Wnt signaling for endoderm differentiation^19^ and motivated by the observation that the Wnt inhibitor Sfrp1 was the most strongly upregulated gene in the AVE cells in BELAs, we explored how manipulating Wnt signaling affected AVE differentiation. Addition of the Wnt agonist Chir99021 completely abrogated OTX2 and Cer1:H2B-Venus expression (Figures 5C-5F), but not GATA6 expression (Figure S4). Inhibition of Wnt signaling with the small molecule XAV939, and inhibition of Wnt secretion with the porcupine inhibitor IWP2 in contrast increased the expression of both Cer1:H2B-Venus and OTX2 (Figures 5C-5F). XAV treatment was more effective than IWP2 treatment and triggered OTX2 expression in almost all cells, while maintaining GATA6 expression (Figures 5C, 5D, 5F, and S4). Thus, the exogenous activation of Activin/Nodal signaling, combined with the inhibition of endogenous Wnt signaling allows the efficient differentiation of AVE cells following forced GATA expression in naïve pluripotent cells. The strong effects of Wnt signaling manipulation on AVE differentiation furthermore suggest that the local inhibition of Wnt signaling through secreted inhibitors contributes to the formation of AVE nests.

## Discussion

Here, we report the differentiation of cohorts of AVE cells in bilayered embryo-like aggregates generated from mouse ESCs. We identify the underlying signaling events between embryonic and extraembryonic cells, and use this knowledge to develop a 2D AVE differentiation protocol. With this protocol, we demonstrate that an antagonism between tissue-intrinsic Wnt signals and Nodal signals coming from both the Epi and the AVE itself control AVE differentiation.

To investigate mechanisms of lineage crosstalk between the Epi and the VE, we used an experimental system where both lineages are established in reproducible proportions from a single starting population through cell-cell communication via FGF4.^9^ This approach contrasts with previous studies, where bilayered aggregates have been formed by mixing ESCs with established XEN cell lines,^20^ by mixing wild-type ESCs with GATA-inducible ESCs,^21^ or by chemical conversion of ESCs towards the VE lineage.^22^ Consistent with our results, these previous studies found that the Epi core induces an embryonic identity in the overlying VE.^22^ They also reported expression of the AVE marker Lefty1 in the VE,^20, 21^ but whether these Lefty1-expressing cells had acquired an AVE identity remained unclear. Our scRNAseq analysis demonstrates that a subset of VE cells in BELAs differentiate into AVE in the absence of an ExE. We speculate that AVE differentiation in BELAs benefits from the specification of the Epi and the VE from a single starting cell population, which closely recapitulates the situation in the embryo.

Using methods to fully direct ESC differentiation towards either Epi or VE, we were able to compare the behavior of pure populations of these lineages with that of mixed populations that form BELAs. In contrast to Epi cells that require exogenous extracellular matrix cues to form cysts,^23^ we find that pure cultures of VE cells spontaneously form cystic structures that resemble the outer layer of BELAs.^24^ This finding suggests that the VE templates the formation of an organized Epi epithelium through the presentation of an extracellular matrix scaffold. Subsequent AVE differentiation in turn is dependent on the presence of the Epi core in BELAs. In line with previous studies from the embryo, we demonstrate that Epi-derived Activin/Nodal signals underlie this inductive event,^4, 25^ thereby representing another swing of a pendulum of interactions between epiblast and VE.^22^

Surprisingly, we find that AVE differentiation occurs in clusters of cells, both in BELAs and in the 2D differentiation protocol where ActivinA is applied globally. Current theories for AVE differentiation in the embryo posit that ExE-derived BMP signals restrict the differentiation of AVE precursors to VE cells at the distal tip of the egg cylinder.^3, 5, 6^ Our observation of restricted AVE differentiation in the absence of an ExE, together with the modest effects of BMP signaling activation and inhibition in the 2D protocol, suggests that this model is incomplete and that other, tissue-intrinsic mechanisms contribute to specifying AVE cells within the bulk of the VE. The strong changes in AVE differentiation upon activating or blocking Wnt signaling with small molecules identify Wnt signaling as an important regulator of AVE differentiation. This idea is further supported by the specific expression of soluble Wnt inhibitors such as Sfrp1 and Sfrp5 in the AVE, as well as impaired AVE precursor differentiation in *Apc^Min/Min^* embryos, where Wnt signaling is hyperactive.^26^ We note that downregulation of Wnt signaling is also required for definitive endoderm differentiation,^19^ a lineage that bears transcriptional similarity to the AVE, thus pointing to a general role of Wnt dynamics in regulating endoderm differentiation.

In addition to its role in endoderm differentiation, Wnt signaling plays a key role in maintaining näive pluripotency in the epiblast of the preimplantation embryo, and its downregulation is required for the transition to rosette-stage pluripotency.^27–29^ Our findings raise the possibility that modulation of Wnt signaling from the VE helps coordinate developmental progression in embryonic and extraembryonic lineages.

Finally, besides regulating developmental progression in the epiblast and the VE in general, tissue-intrinsic Wnt signals may contribute to the patterned differentiation of AVE cells in local clusters. Wnt-based patterning mechanisms underlie hair follicle differentiation in the mouse skin, and axis formation during planarian regeneration.^30, 31^ Further studies using BELAs and the 2D AVE differentiation system will be required to identify the components and the topology of the cell-cell communication network for AVE differentiation and patterning. Such mechanisms that do not rely on an external BMP gradient may explain axis formation in disc-shaped non-rodent embryos.

## Acknowledgements

We thank M. Sandhaus for his contributions to the early stages of this project, and Sarah Teckhaus for help with generating Nodal mutant cells. We are grateful to Shankar Srinivas and Antonia Scialdone for sharing unpublished sequencing data. We thank P. Bastiaens, current and former members of the Schröter group, and all members of the Department for Systemic Cell Biology for support, stimulating discussions and conceptual input on the project, as well as Alfonso Martinez Arias, Nicholas Rivron, Stefan Semrau, Naomi Moris, Vikas Trivedi for feedback on earlier versions of this manuscript. This work was supported by the VW foundation (project no. A130140 “OntoTime”), the German Research foundation (project no. 441798639), the German Center for the protection of laboratory animals (Bf3R project no. 1328-567), an ERC consolidator grant (MORPHEUS, 101043753 to I.B.) and the Max Planck Society.

## Author contributions

Conceptualization and Methodology: S.S., C.S.; Validation: S.S., M.M.; Formal analysis: S.S.; M.F.; Investigation: S.S., M.F., M.M., Y.S.K., C.S.; Data Curation: M.F.; Writing - Original Draft: S.S., M.F., C.S.; Writing - Review and Editing: All authors; Visualization: S.S., M.F.; Supervision: I.B., C.S.; Funding Acquisition: I.B., C.S.

## Materials and Methods

### Cell lines

All cell lines used in this study were on an E14tg2a background.^32^ The inducible Tet::GATA4-mCherry (iGATA) lines have previously been described.^9^ We used two different clones in this study, C5 and C6, that differ in their induction rate. Cells were maintained on fibronectin coated dishes in N2B27-based medium supplemented with 1 µM PD0325901 (SeleckChem), 10 ng/ml LIF (protein expression facility, MPI Dortmund), and 3 µM CHIR99021 (Tocris), referred to as 2i + LIF.^33^ N2B27 was prepared as a 1:1 mixture of DMEM/F12 and Neuropan Basal Medium (both from PAN Biotech), supplemented with 1X N2 and 1X B27 supplements, 1X L-Glutamax, 0.0025% BSA, and 0.2 mM ß-mercaptoethanol (all from ThermoFisher). All iGATA4 cell lines were kept under constant selection with 200 µg/ml G418 (Sigma) to prevent silencing of the inducible transgene. Cells were cultured at 37°C with 5% CO_2_, and regularly tested for mycoplasma contamination.

### Mouse strains

Mice used for tetraploid complementation were of the B6C3F1 or CD1 strains and were raised in-house.

### Generation of mutant and transgenic ESC lines

To generate a Cer1 reporter in the iGATA cell line, the Cer1 promoter region 4kb upstream of the start codon was amplified from genomic DNA, a puromycin resistance cassette and a H2B-Venus sequence were amplified from Sprouty4 targeting vectors described in Morgani et al., 2018,^34^ and Raina et al., 2021.^9^ All three fragments were cloned via Gibson assembly using a HiFi DNA assembly kit (NEB) into a vector backbone containing PiggyBac transposition sites.^35^ The Cer1:H2B-Venus reporter construct was co-transfected with CAG-pBASE^35^ using Lipofectamine 2000 (ThermoFisher) according to manufacturer’s instructions. Cells were selected with 1.5 µg/ml puromycin (Sigma) starting 24 hours after transfection. Colonies were picked one week after transfection, expanded and evaluated for co-localization of Cer1 reporter activity and *Cer1* mRNA.

CRISPR/Cas9 was used to mutate the *Nodal* locus in iGATA ESCs (clone C6) and one subclone carrying the Cer1:H2B-Venus reporter construct. sgRNAs 5’-CCCCATGGACATACCCACTG-3’ and 5’-CCAGTCGAGCAGAAAAGTGT-3’ defining a 244 bp region in *Nodal* exon 2 were cloned into pX458 (Addgene plasmid #48138) or pX459 (Addgene plasmid #48139) using BbsI (NEB) according to Ran et al., 2013.^36^ Cells were transfected using Lipofectamine 2000 (Thermo Fisher Scientific) according to manufacturer’s instructions. To enrich for transfectants, cells were either selected with 1.5 µg/ml puromycin for two days, or flow sorted for GFP-expression before seeding at clonal density. We established several clonal lines, and used primers 5’-GTGGACGTGACCGGACAGAACTG-3’ and 5’-GGCATGGTTGGTAGGATGAAACTCC-3’ to PCR-amplify a sequence around the CRISPR mutation site. Clones that gave a shortened amplicon compared to the wild type were chosen for further analysis, and the exact sequence of the mutated alleles was determined by Sanger sequencing.

To generate constitutively labeled cell lines, we modified a piggybac vector for the constitutive expression of H2B-Cerulean^37^ by either replacing its puromycin resistance cassette with a blasticidin resistance cassette from pCX-H2B-Cerulean-IRES-bsd^38^ using restriction enzymes PmiI and and PstI, or by replacing the H2B-Cerulean sequence with an mCherry coding sequence using restriction enzymes SpeI and NotI. Vectors were co-transfected with CAG-pBASE^35^ using Lipofectamine 2000 according to the manufacturer’s instructions, and transfected cells were selected with 15 µg/ml blasticidin 48 h after transfection. Four days after transfection, cells were flow sorted for the expression of fluorescent proteins, and seeded a clonal density. Several clones were expanded, and two to three suitable clones with homogeneous, moderate H2B-Cerulean and/or mCherry fluorescence were selected by epifluorescence microscopy for further experiments.

### Differentiation of pure cultures of PrE cells and subsequent AVE differentiation

Pure cultures of PrE cells from iGATA clone C6 were obtained by inducing with 0.5 µg/ml doxycycline in 2i + LIF for 8 hours, followed by another 16 hours of doxycycline treatment in N2B27 supplemented with 10 ng/ml FGF4 and 1 µg/ml heparin. To obtain these cultures from iGATA clone C5, a 4 hours pulse with 0.5 µg/ml doxycycline in 2i + LIF followed by further culture in N2B27 supplemented with FGF4 and heparin was sufficient. Clone C5 was used for the data shown in Figure S1B, in all other instances, clone C6 was used.

To differentiate AVE cells from these cultures, cells were additionally treated with 50 ng/ml ActivinA upon media change from 2i + LIF to N2B27. Approximately 24 hours after the start of doxycycline treatment, cells were re-seeded at a total density of 25,000 to 30,000 cells/cm^2^ on fibronectin-coated dishes and cultured for three days in N2B27 supplemented with 10 ng/ml FGF4, 1 µg/ml heparin and 50 ng/ml ActivinA.

### Formation of BELAs, VE- and Epi-cysts

BELAs were generated by inducing iGATA ESCs with 0.5 µg/ml doxycycline in 2i + LIF for 8 hours, followed by a media change to N2B27 for 16 hours. Cells were then seeded at a density of 30,000 cells/cm^2^ on dishes that had been coated with 0.1% gelatin in PBS for 30 minutes. Floating aggregates were collected for further analysis at indicated time points.

VE cysts were generated from pure cultures of PrE cells differentiated as described above, followed by re-seeding onto gelatin-coated dishes at a density of 30,000 cells/cm^2^ in N2B27 medium supplemented with 10 ng/ml FGF4 and 1 µg/ml heparin.

Cysts of Epi cells were made according to Bedzhov and Zernicka-Goetz, 2014,^23^ with minor modifications. iGATA ESCs were detached, resuspended in growth factor-reduced matrigel (Corning) and plated as 25 µl drops on µ-slides (ibidi). The slides were incubated at 37°C to allow the matrigel to solidify and then filled with prewarmed N2B27 or 2i + LIF medium.

### Generation of epiblast-specific Nodal knock-out embryos

Epiblast-specific Nodal knock-out embryos were generated via tetraploid complementation. Donor embryos used for tetraploid complementation were derived from the B6C3F1 strain and foster mothers for embryo transfer experiments were from the CD1 background. Briefly, tetraploid morulae were aggregated with Nodal-mutant ESCs^4, 37^ or wild-type E14 ESCs. The aggregated embryos were cultured in KSOM (Millipore) for additional three days, which were then transferred into the uterus of foster mothers. Post-implantation embryonic day (E) 5.5 tetraploid embryos were recovered by manually dissecting the uterus.

### Immunostaining

BELAs and VE cysts in suspension were fixed with 4% paraformaldehyde at room temperature for 1 hour, washed 5 times with phosphate buffered saline (PBS) for 5 minutes each, and then incubated in PBS supplemented with 1% BSA and 0.1% Triton X-100 (PBT+BSA) for 3 hours at room temperature, followed by incubation with primary antibodies diluted in PBT+BSA at 4°C overnight. Primary antibodies used were anti-Oct3/4 (POU5F1, Santa Cruz Biotechnology sc-5279 1:100), anti-E-Cadherin (CDH1, Takara M108, 1:200), anti-GATA6 (R&D, RF1700, 1:200), anti-CD29 (ITGB1, BD Pharmingen 562153, 1:100), anti-LAM (Sigma L9393, 1:750), anti-OTX2 (Neuromics GT15095, 1:200), anti-pERM (Cell Signaling Technology #3141, 1:200), anti-PODXL (R&D MAB1556, 1:200), anti-SOX17 (R&D AF1924, 1:200), anti-ZO-1 (Invitrogen 61-7300, 1:100), and anti-GFP (Abcam ab13970 1:200).

To remove the primary antibody solution, the aggregates were washed five times with PBT+BSA. Aggregates were then incubated overnight at 4°C with secondary antibodies diluted in PBT+BSA. Secondary antibodies from Invitrogen/Life Technologies were Alexa Fluor-conjugated and used at 4 µg/ml. Nuclei were stained with Hoechst 33342 dye at 1 µg/ml (Invitrogen). The secondary antibody solution was removed by 5 washes with PBS supplemented with 0.1% Triton X-100. The aggregates were resuspended in PBS and mounted onto µ-slides (ibidi).

Epi cysts in matrigel and cells grown in µ-slides (ibidi) were stained similarly, but with extended incubation and wash times for Epi cysts, and shortened times for cells grown as 2D layers. Samples were mounted in mounting solution consisting of 16% PBS, 80% glycerol, and 4% n-propyl-gallate.

Post-implantation embryos were fixed in 4% PFA for 20 min, and washed twice in wash buffer containing 1% fetal calf serum (FCS) in PBS. The embryos were then permeabilized in 0.1 M glycin/0.3% Triton-X in PBS for 10 min, and washed twice in the wash buffer. The embryos were then incubated with primary antibody anti-CER1 (R&D, MAB1986) in blocking buffer containing 2% FCS in PBS overnight at 4°C. After two washes in wash buffer, embryos were incubated with secondary antibodies and DAPI in blocking buffer, which were washed twice on the next day. The stained embryos were mounted in droplets of wash buffer on 35 mm µ-Dish glass bottom plates (ibidi), covered with mineral oil and stored at 4°C until imaging.

### In situ HCR

For third generation in situ HCR we used probe sets, wash and hybridization buffers together with corresponding Alexa Fluor-labeled amplifiers from Molecular Instruments.^39^ Staining was performed according to the manufacturer’s instructions. Briefly, samples were fixed for 15 min to 1 h with 4% paraformaldehyde, washed four times with PBS with 0.1% Tween 20 (PBST) and permeabilized at least overnight in 70% ethanol at -20°C. Samples were then washed twice with PBST, and equilibrated in probe hybridization buffer for 30 min at 37°C. Transcript-specific probes for *Otx2* (NM_144841.5)*, Gata6* (NM_010258) and *Cer1* (NM_009887.2) were designed by Molecular Instruments. Probes were used at a final concentration of 4 nM in probe hybridization buffer and incubated overnight at 37°C. To remove the probe solution, the sample was washed four times with probe wash buffer preheated to 37°C and once with 5x SSC with 0.1% Tween 20 (SSCT). Samples were then equilibrated in amplification buffer for 30 min at room temperature. Alexa Fluor-labeled amplifiers were used at a final concentration of 60 nM together with Hoechst 33342 dye at 1 µg/ml and incubated overnight at room temperature. The amplifier solution was removed by six washes with 5x SSCT. Stained BELAs were resuspended in PBS and mounted on an ibidi µ-slide for imaging. 2D cultures were mounted in mounting solution consisting of 16% PBS, 80% glycerol, and 4% n-propyl-gallate.

### Imaging

Cells for long-term imaging (Figures 1B and S1B) were seeded at a density of 30,000 cells/cm^2^ on 6-well plates (Sarstedt) or 8-well µ-slides plates (ibidi) and allowed to attach for 1-2 hours before the start of imaging. Time-lapse movies were recorded with a 20x 0.5 NA air objective on an Olympus IX81 widefield microscope equipped with a stage top incubator (ibidi), LED illumination (pE4000, CoolLED) and a c9100-13 EMCCD (Hamamatsu) camera. Hardware was controlled by MicroManager software,^40^ and tile scans were stitched in FIJI using the pairwise stitching plugin.^41^ Live VE cysts in Figure 1I, J were imaged on a Leica DM IRB widefield microscope using a 20x 0.4 NA (Figure 1I) or a 40x 0.55 NA (Figure 1J) phase contrast objective. Stained BELAs, embryos, and stained cells in 2D culture were imaged on a Leica SP8 confocal microscope (Leica Microsystems) with a 63x 1.4 NA oil immersion objective. Cultures to determine the clonal composition of AVE clusters in 2D culture (Figure 5B) were fixed, incubated with SYTO Deep Red Nucleic Acid Stain (ThermoFisher) for one hour, and imaged with a 20x 0.5 NA air objective on an Olympus IX81 widefield microscope equipped with LED illumination (pE4000, CoolLED) and an iXon 888 EM-CCD camera (Andor). Hardware was controlled by MicroManager software^40^ and tile scans were stitched in FIJI using the pairwise stitching plugin.^41^

For light sheet imaging, fixed and stained aggregates were resuspended in low melting agarose and placed in 1.5 mm U-shaped capillaries (Leica). Capillaries were placed into water filled 35mm high glass bottom µ-dishes (ibidi). Images were acquired using an HC Fluotar L 25x 0.95 NA water DLS TwinFlect 2.5 mm detection objective and an HC PL Fluotar 5x 0.15 NA illumination objective on a Leica TCS SP8 digital light sheet microscope. 3D animations were created using the Leica X application suite. Z-Stack images were processed and quantified using FIJI and Imaris.

### Flow cytometry

Cells for flow cytometry were detached from culture vessels, fixed in 4% paraformaldehyde for 15 min, washed with PBS and then incubated in PBS + 1% BSA + 0.25% Saponin (PBSap) for 30 min at room temperature. Afterwards, cells were incubated with primary antibodies diluted in PBSap at 4°C overnight. The next day, cells were washed three times in PBSap and incubated with secondary antibodies diluted in PBSap for at least one hour. Cells were washed three times in PBSap, and passed through a cell strainer and analyzed immediately on a LSRII flow cytometer (BD Biosciences). Live cells were sorted on a FACS Aria Fusion (BD Biosciences). Flow cytometry data was analyzed with FlowJo (BD Biosciences).

### ScRNAseq sample preparation

BELAs and VE cysts were generated as described above. Between 100 and 200 BELAs and VE cysts were manually picked under a dissection microscope for further processing. We selected round aggregates and cysts, and excluded structures that contained a large number of dead cells, or that were unusually big or small. For the VE cysts, we also aimed at excluding structures that contained a clearly visible core of putative Epi-like cells, which likely arise from insufficiently induced cells. Both BELAs and VE cysts were gently spun down, resuspended in 1 ml Accutase and incubated at 37°C for 10 min, followed by mechanical dissociation by pipetting and further incubation in Accutase for 5 min. Next, cells were spun down, washed in PBS, and resuspended in a small volume of PBS + 0.5% BSA.

To generate Epi cysts for RNA sequencing, single cells were seeded in matrigel and cultured in 2i + LIF for one day. Then, medium was changed to N2B27, and cells were cultured for another 3 days. Cysts were recovered from matrigel by incubation in recovery solution (Corning) for 20 min on ice. Next, cysts were gently spun down and dissociated with Accutase as described for BELAs and VE cysts above. To remove residual matrigel, dissociated cells were washed once with recovery solution and twice with ice-cold PBS, followed by resuspending in a small volume of PBS + 0.5% BSA.

### ScRNAseq library preparation and sequencing

Cells from all three samples were counted, and each sample was mixed with H_2_O and RT master mix from the Chromium Next GEM Single Cell 3’ GEM, Library & Gel Bead Kit v3.1 (10x Genomics) to obtain a cell density required for targeting 1000 (Epi and VE cysts) or 2000 (BELAs) cells. Cell suspensions were loaded on a Chromium Controller (10x Genomics) to partition cells with gel beads in emulsion. Reverse transcription, cDNA recovery and amplification, and sequencing library construction were performed according to manufacturer’s instructions (10x Genomics ChromiumNextGEMSingleCell_v3.1_Rev_D). We chose 12 PCR cycles for cDNA amplification, and 13 PCR cycles for index PCR. Concentration and insert size of sequencing libraries were determined with a BioAnalyzer High Sensitivity DNA Assay (Agilent). Libraries were sequenced by paired-end Illumina sequencing on a NovaSeq6000 instrument with a read length of 150 bp. We first performed sequencing at shallow depth with a target of 10.000 reads per cell, to confirm capturing of an appropriate number of high-quality single-cell transcriptomes. Subsequently, deeper sequencing was performed, to obtain between 100.000 and 150.000 reads per cell.

### ScRNAseq data analysis

Demultiplexing, alignment to the mouse genome mm10 (GENCODE vM23/Ensembl 98, from 10x Genomics) and read quantification was performed with CellRanger (10x Genomics, v4.0.0). Subsequent analysis was carried out in R using Seurat v4.1.1.^42^ We first filtered out cells with less than 4000 different features detected and with more than 10% of the reads mapped to mitochondrial genes. SCTransform^42^ was used to normalize and scale the molecular count data. For Uniform Manifold Approximation and Projection (UMAP) representation and clustering, shared cell populations were matched across samples using Seurat’s integration algorithm for SCTransformed data with reciprocal PCA to identify anchors. Differentially expressed genes between the clusters resulting from Louvain clustering were identified with the FindMarker function based on the SCTransform normalized data, and sorted by fold-change.

ScRNAseq data from the developing mouse embryo was obtained from two publications: Raw counts of the E3.5 to E8.75 embryo dataset from Nowotschin et al., 2019,^13^ including cell type annotations were downloaded from https://endoderm-explorer.com. For visualization, we did not differentiate between the different types of gut tube cells annotated by Nowotschin et al., 2019,^13^ but used “gut tube” as a single label for all these cells. Similarly, we did not differentiate between different samples collected from E8.75 embryos, but pooled these groups with a single E8.75 label. This dataset was integrated with all single-cell transcriptomes from our study in SCANPY,^43^ using log1p-transformed counts after normalization of our data to 10000 reads per cell. The asymmetric integration and label transfer was performed with ingest and cell type proportions were visualized in R using a custom heatmap function based on pheatmap.

ScRNAseq data and annotations of an embryo dataset focused on AVE development was obtained from the authors.^16^ Integration of this dataset was performed with BELA cells from clusters 3 and 4 in Figure 3A only, using the same pipeline as for the Nowotschin dataset.

### Cell-Cell communication analysis

For the inference of cell-cell communication events from scRNAseq data we used LIANA, a LIgand-receptor ANalysis frAmework.^18^ To identify cell-cell communication events in BELAs, we only used transcriptomes from this sample, and grouped them into two lineages according to the clustering in Figure 3A: All cells from clusters 1 and 2 were grouped as Epi, and cells from clusters 3 and 4 were grouped as VE. The consensus database for ligand-receptor interactions was matched to its mouse ortholog genes using the omnipath database, and interactions were ordered by their consensus rank obtained from LIANA. For Figure 4A, the top 20 interactions were displayed as an undirected adjacency graph.

### Quantification and statistical analysis

Quantitative data are represented as mean ± SD. Number of repeat experiments is stated in figure legends, with N indicating number of biological replicates, and n indicating number of independent samples within an experiment. For flow cytometry experiments in Figures 4, 5, S5, and S6, at least n = 20.000 cells were analyzed for each condition in each biological replicate. Statistical analysis was performed in GraphPad Prism 8 (v8.4.3), using unpaired or paired ratio t-tests as indicated in the figure legends. * and ** indicate p ≤ 0.05 and p ≤ 0.005, respectively. Significance of differential gene expression between clusters in scRNAseq data was assessed with a Wilcox likelihood-ratio test in R.

### Data and code availability

Single-cell RNA-sequencing data generated in this study has been deposited at the NCBI gene expression omnibus repository under accession number GSE198780. All code used for analysis and visualization, together with a list of the R packages used, is available on GitHub at https://github.com/Schroeterlab/BELAs_Schumacher_et_al. Any additional information required to reanalyze the data reported in this paper is available from the authors upon request.

## Supplementary Materials

## Supplementary videos

**Video S1. Time-lapse imaging of BELA formation, related to Figure 1**

Mixtures Epi and PrE cells were differentiated from a single culture and re-seeded in N2B27 medium on gelatin-coated dishes 16 h after the end of the doxycycline pulse. Scale bar: 200 μm, frame rate: 2 images/h.

**Video S2**. **Light sheet imaging of BELA stained for GATA6 and POU5F1, related to** Figure 1

3D rendering and animation of a BELA stained for POU5F1 (green) and GATA6 (magenta) and imaged by light sheet microscopy.

**Video S3**. **Organization of basement membrane in a BELA detected by light sheet imaging, related to** Figure 1

Animation of Z-stack of same BELA as in Video S2, but now also showing Laminin staining in yellow.

**Video S4**. **Light sheet imaging of Cer1:H2B-Venus expression in a BELA, related to** Figure 3

3D rendering and animation of a BELA made from Cer1:H2B-Venus reporter cells stained for POU5F1 (green) and Cer1:H2B-Venus (yellow), and imaged by light sheet microscopy.

## Supplementary Tables

**Table S1. List of differentially expressed genes between clusters 3 and 4 identified in Figure 3**

**Table S2. Output of LIANA analysis, related to Figure 4**

Table lists potential ligand-receptor interactions between Epi cells (clusters 1 and 2 in Figure 3A), and VE cells (clusters 3 and 4 in Figure 3A) from the BELA sample.

**Figure S1.**
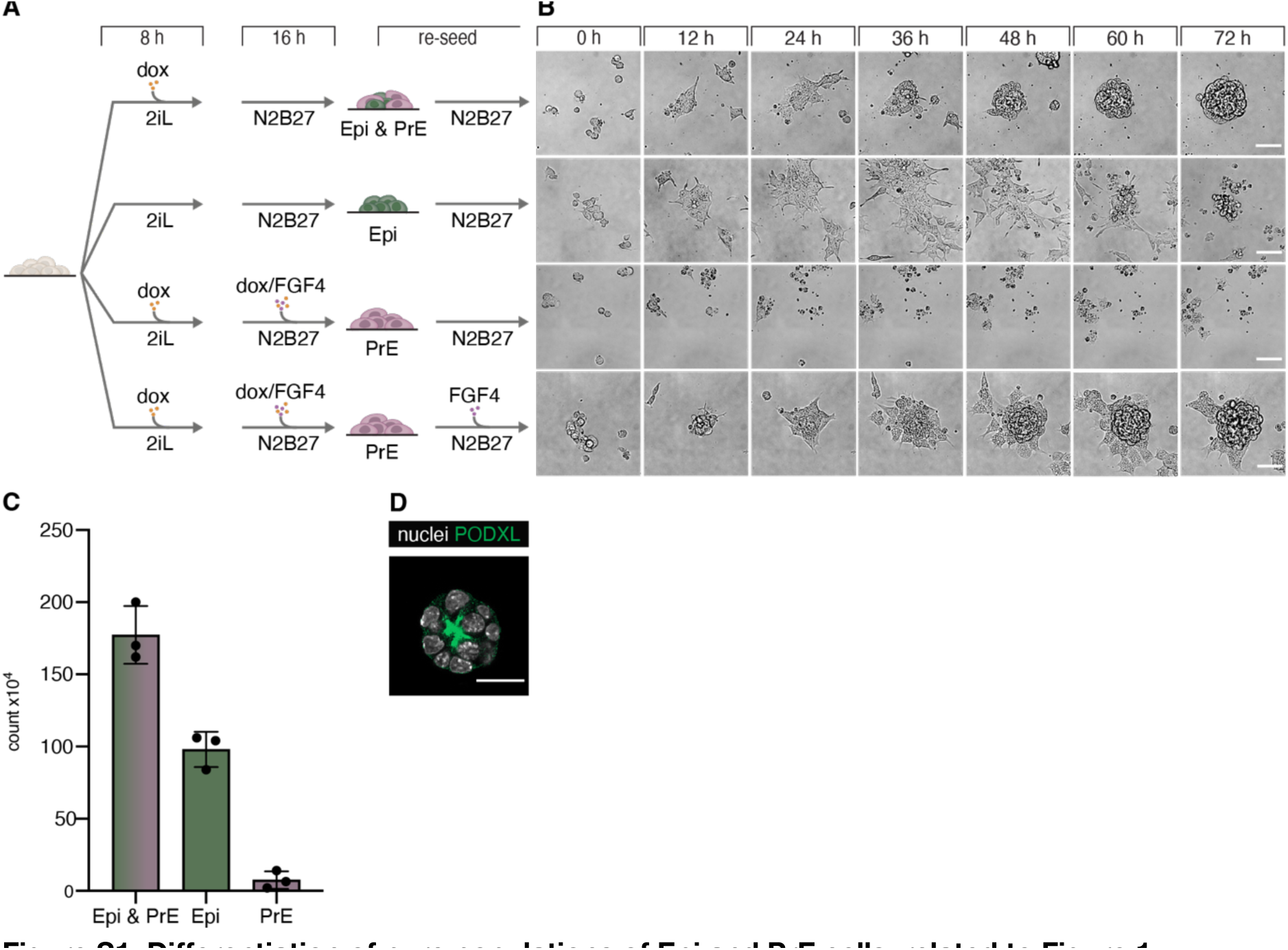
Differentiation of pure populations of Epi and PrE cells, related to Figure 1. (A) Experimental approach to differentiate pure populations of Epi and PrE cells. Top row indicates method to generate mixed cultures of Epi and PrE cells for comparison. (B) Stills from movies of the cell types differentiated as in (A) after re-seeding in N2B27. (C) Quantification of live cells differentiated as in (A) without addition of exogenous FGF4 (first three conditions in (A)) three days after re-seeding. (D) Immunostaining for the polarisation marker PODXL in mESCs seeded in Matrigel and cultured for 2 days in N2B27. Scale bars: 25 µm.

**Figure S2.**
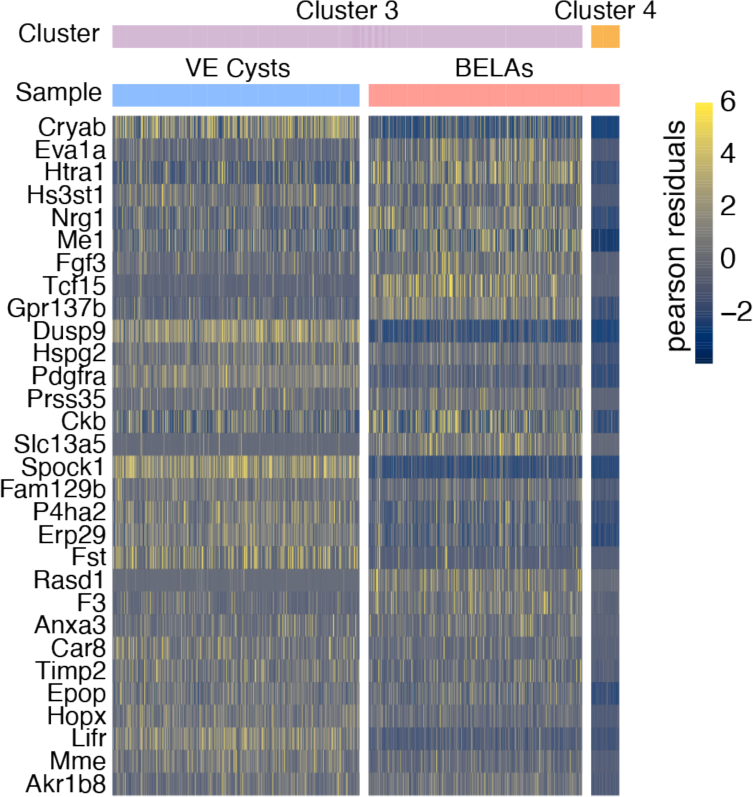
Down-regulated genes in cluster 4 from Figure 3A. Heatmap showing the 30 most down-regulated genes between the cells of cluster 3 and cluster 4 from Figure 3A, ordered by log2-fold change. Single-cell expression is shown as the Pearson residual of the normalized counts.

**Figure S3.**
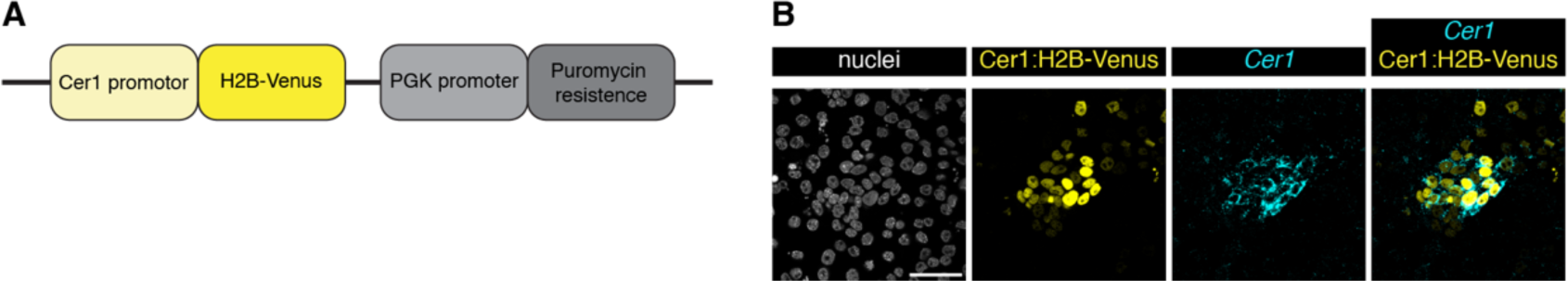
Design and validation of a Cer1:H2B-Venus reporter construct, related to Figure 3. (A) Schematic of the Cer1:H2B-Venus reporter construct. A 4-kb regulatory region of the Cer1 gene^17^ was fused to an H2B-Venus reporter, coupled to a puromycin resistance cassette, and integrated into inducible cells via piggybac transgenesis, (B) Co-expression of H2B-Venus protein (yellow) and Cer1 mRNA (cyan) stained by in situ HCR. Shown is a single confocal section of Cer1:H2B-reporter cells differentiated towards PrE and treated with 50 ng/ml Activin A after removal of 2i medium. Nuclei are labeled with Hoechst33342. Scale bar: 50 µm.

**Figure S4.**
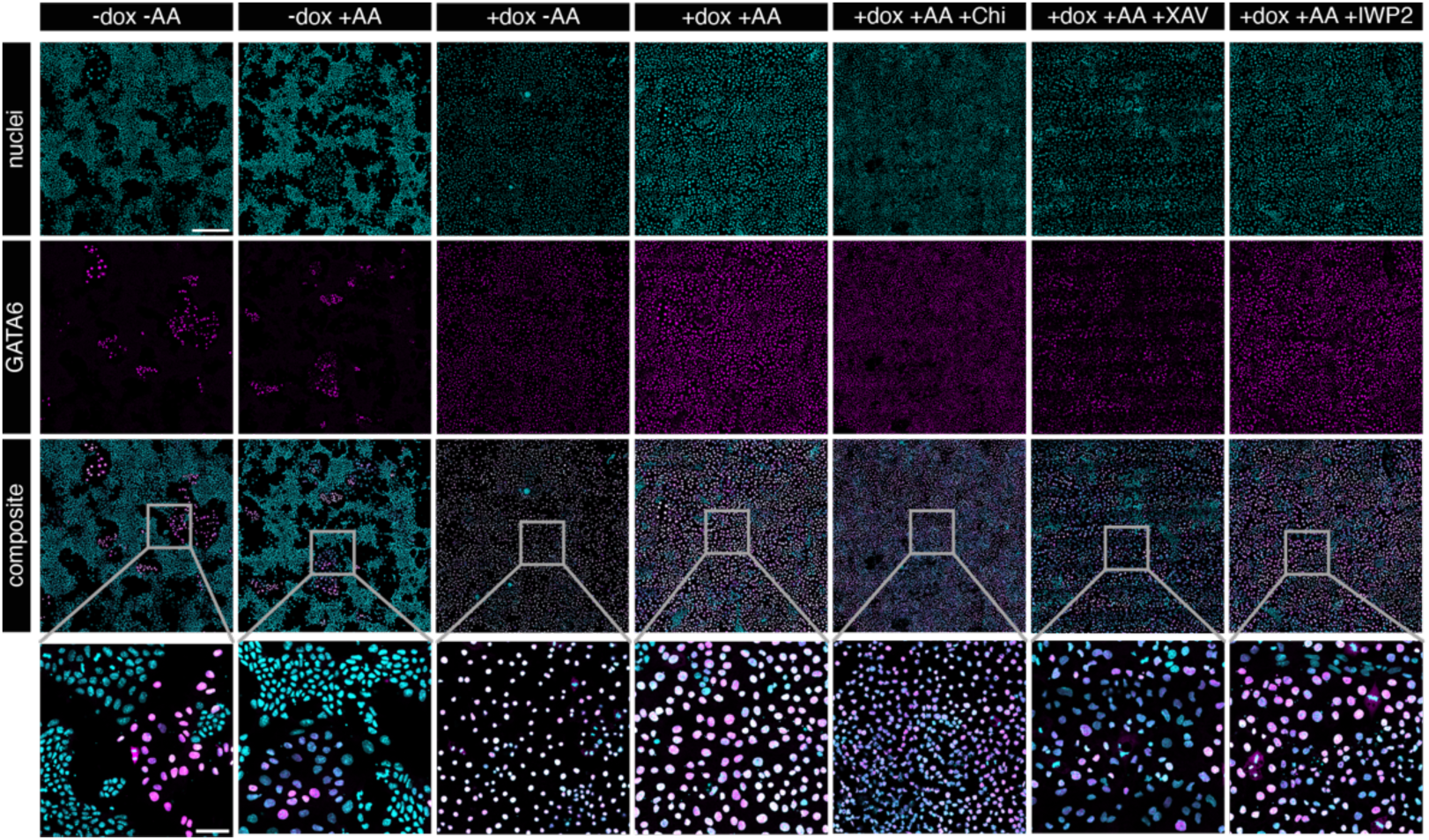
Expression of the endoderm marker GATA6 is dependent on doxycycline treatment of inducible cells, but independent from ActivinA and Wnt signaling, related to Figures 4 and 5. Immunostaining of GATA4-inducible cells cultured for 3 days in the indicated media conditions without (first two columns) or with prior doxycycline induction. GATA6 expression in magenta, nuclei stained with Hoechst33342 shown in cyan. ActivinA was used at a concentration of 50 ng/ml, concentrations of all other supplements were the same as in Figure 5. Note that a similar, low number of GATA6-positive cells is obtained in the absence of doxycycline induction both with and without ActivinA treatment, suggesting that these are a consequence of leaky transgene expression. Scale bars: 250 µm (upper panels); 50 µm (bottom panels).

**Figure S5.**
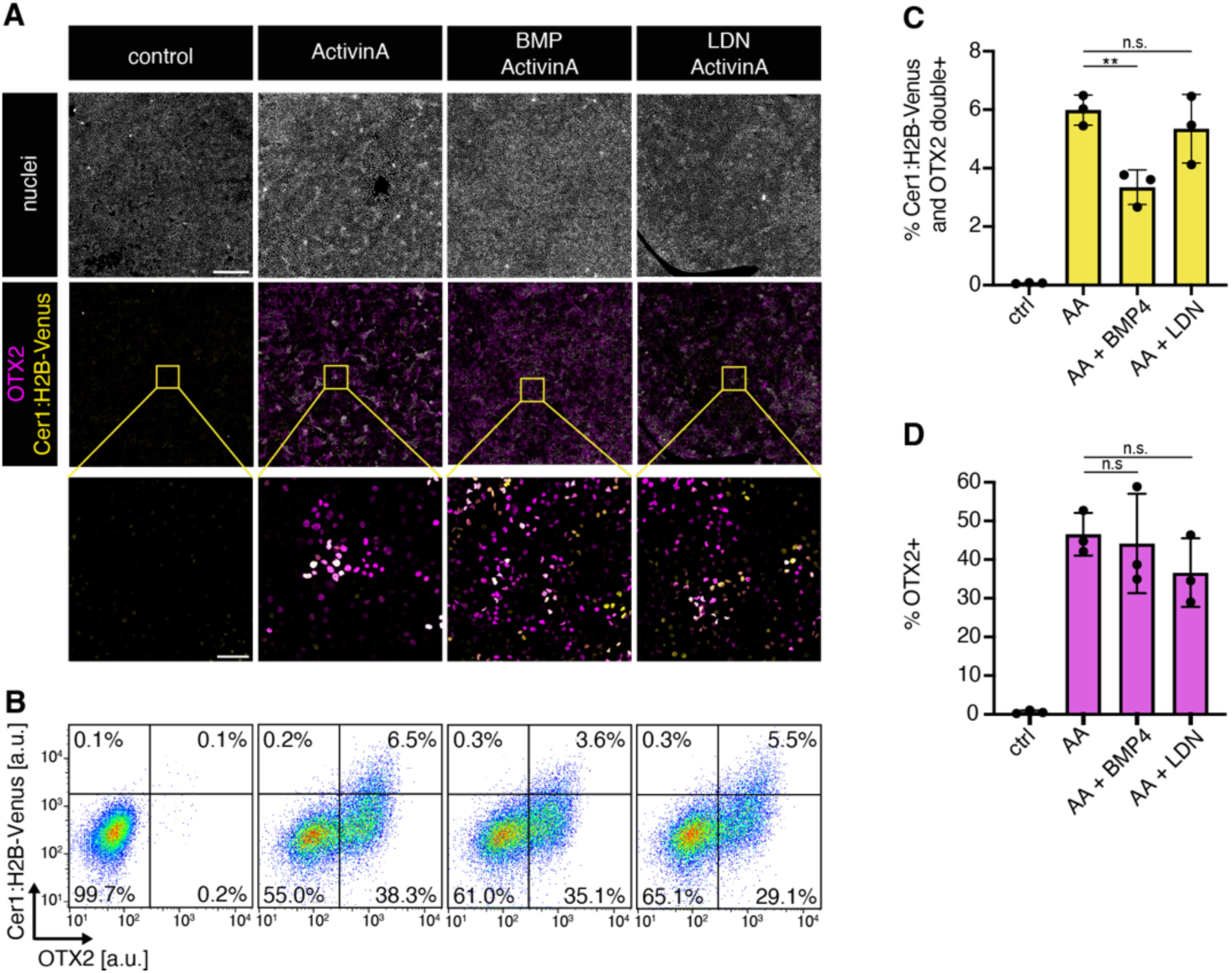
Influence of BMP signaling manipulation on AVE differentiation, related to Figure 5. (A) Immunostaining for OTX2 (magenta) and H2B-Venus (yellow) of Cer1:H2B-Venus reporter cells differentiated for 3 days after an extended doxycycline pulse with 50 ng/ml ActivinA (AA), together with 50 ng/ml BMP4 or 100 nM LDN193189 as indicated. (B) Flow cytometry of cells differentiated and stained as in (C). (C) Mean percentage of Cer1:H2B-Venus; OTX2 double-positive cells differentiated as in (B). N = 3, error bars indicate SD. (D) Same as (C) but showing percentage of OTX2-positive cells. Data for conditions without ActivinA, and with ActivinA but without BMP signaling manipulation are the same as in Figure 5. Scale bars: 500 µm ((A) overview); 50 µm ((A) inset). ** in (C) indicates p ≤ 0.005, n.s. indicates p > 0.05 determined by a two-tailed, unpaired t-test.

**Figure S6.**
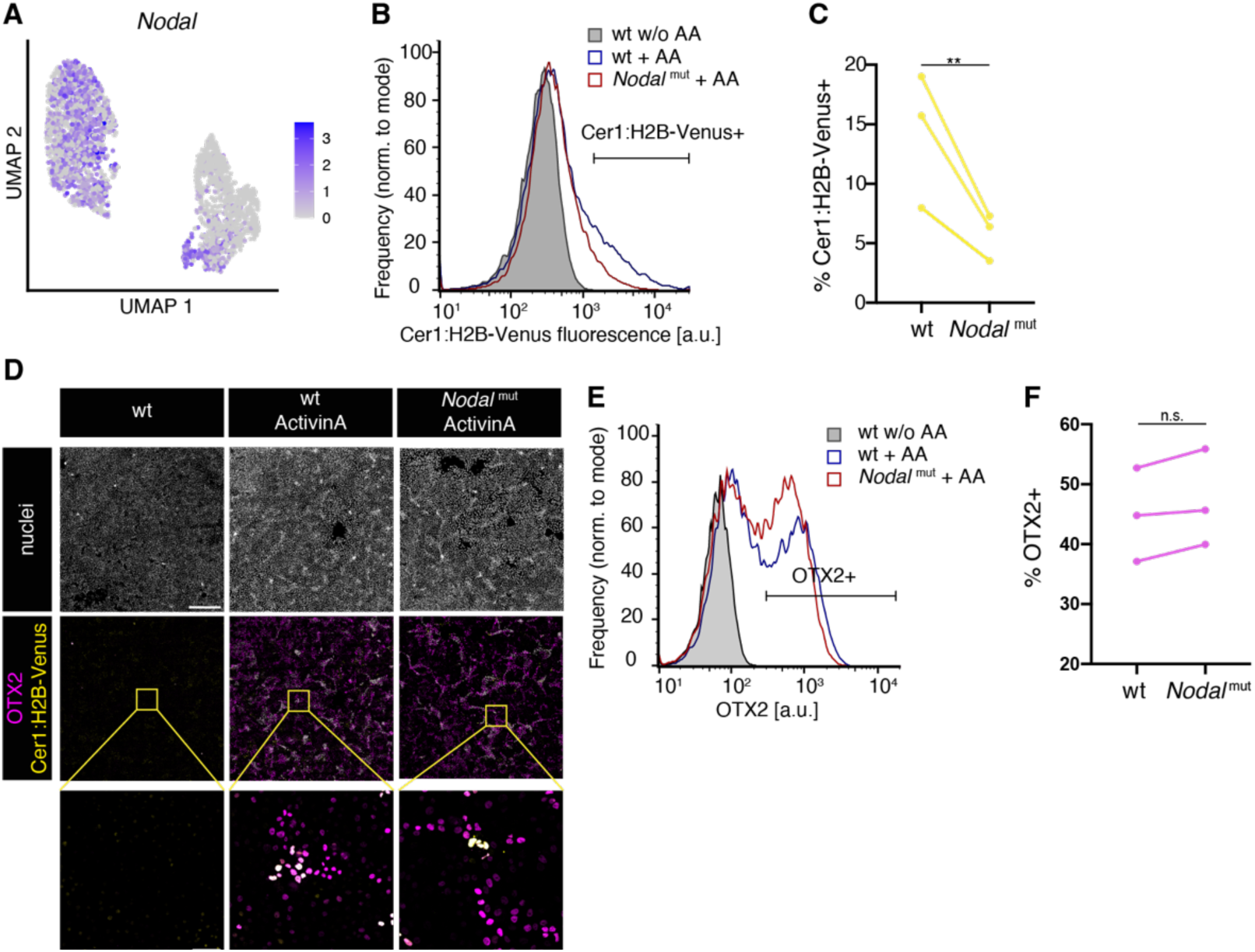
AVE differentiation in Nodal-mutant cells, related to Figure 5. (A) Nodal expression from single-cell sequencing data, shown on UMAP plot from Figure 2B. (B) Flow cytometry of wild-type and Nodal-mutant Cer1:H2B-Venus reporter cells differentiated for 3 days after an extended doxycycline pulse with 50 ng/ml ActivinA. (C) Percentage of Cer1:H2B-Venus-positive cells differentiated as in (B) from N = 3 independent experiments. (D) Immunostaining for OTX2 (magenta) and H2B-Venus (yellow) of wild-type and Nodal-mutant Cer1:H2B-Venus reporter cells as in (B). (E) Flow cytometry of OTX2 staining of cells differentiated as in (B). (F) Mean percentage of OTX2-positive cells differentiated as in (B) from N = 3 independent experiments. Variability in background staining intensities precluded comparison of Cer1:H2B-Venus signal in fixed and stained cells in (D) - (F). Data from wild-type cells with and without ActivinA are the same as in Figure 5. Scale bars: 500 µm ((D) overview); 50 µm ((D) inset). ** in (C) indicates p ≤ 0.005, n.s. indicates p > 0.05 determined by a two-tailed, paired ratio t-test.

## References

1. Hoshino, H., Shioi, G., and Aizawa, S. (2015). AVE protein expression and visceral endoderm cell behavior during anterior–posterior axis formation in mouse embryos: Asymmetry in OTX2 and DKK1 expression. Dev. Biol. 402, 175–191. 10.1016/J.YDBIO.2015.03.023.

2. Nowotschin, S., Costello, I., Piliszek, A., Kwon, G.S., Mao, C., Klein, W.H., Robertson, E.J., and Hadjantonakis, A.-K. (2013). The T-box transcription factor Eomesodermin is essential for AVE induction in the mouse embryo. Genes Dev. 27, 997–1002. 10.1101/gad.215152.113.

3. Stower, M.J., and Srinivas, S. (2018). The Head’s Tale: Anterior-Posterior Axis Formation in the Mouse Embryo. Curr. Top. Dev. Biol. 128, 365–390. 10.1016/bs.ctdb.2017.11.003.

4. Brennan, J., Lu, C.C., Norris, D.P., Rodriguez, T.A., Beddington, R.S.P., and Robertson, E.J. (2001). Nodal signalling in the epiblast patterns the early mouse embryo. Nature 411, 965–969. 10.1038/35082103.

5. Rodriguez, T.A., Srinivas, S., Clements, M.P., Smith, J.C., and Beddington, R.S.P. (2005). Induction and migration of the anterior visceral endoderm is regulated by the extra-embryonic ectoderm. Development 132, 2513–2520. 10.1242/dev.01847.

6. Yamamoto, M., Beppu, H., Takaoka, K., Meno, C., Li, E., Miyazono, K., and Hamada, H. (2009). Antagonism between Smad1 and Smad2 signaling determines the site of distal visceral endoderm formation in the mouse embryo. J. Cell Biol. 184, 323–334. 10.1083/jcb.200808044.

7. Ma, H., Zhai, J., Wan, H., Jiang, X., Wang, X., Wang, L., Xiang, Y., He, X., Zhao, Z.-A., Zhao, B., et al. (2019). In vitro culture of cynomolgus monkey embryos beyond early gastrulation. Science (80-.). 366. 10.1126/science.aax7890.

8. Molè, M.A., Coorens, T.H.H., Shahbazi, M.N., Weberling, A., Weatherbee, B.A.T., Gantner, C.W., Sancho-Serra, C., Richardson, L., Drinkwater, A., Syed, N., et al. (2021). A single cell characterisation of human embryogenesis identifies pluripotency transitions and putative anterior hypoblast centre. Nat. Commun. 12, 3679. 10.1038/s41467-021-23758-w.

9. Raina, D., Bahadori, A., Stanoev, A., Protzek, M., Koseska, A., and Schroter, C. (2021). Cell-cell communication through FGF4 generates and maintains robust proportions of differentiated cell types in embryonic stem cells. Dev. 148, dev199926. 10.1242/dev.199926.

10. Wamaitha, S.E., del Valle, I., Cho, L.T.Y., Wei, Y., Fogarty, N.M.E., Blakeley, P., Sherwood, R.I., Ji, H., and Niakan, K.K. (2015). Gata6 potently initiates reprograming of pluripotent and differentiated cells to extraembryonic endoderm stem cells. Genes Dev. 29, 1239–1255. 10.1101/gad.257071.114.

11. Hermitte, S., and Chazaud, C. (2014). Primitive endoderm differentiation: From specification to epithelium formation. Philos. Trans. R. Soc. B Biol. Sci. 369, 20130537. 10.1098/rstb.2013.0537.

12. Saiz, N., and Plusa, B. (2013). Early cell fate decisions in the mouse embryo. REPRODUCTION 145, R65–R80. 10.1530/REP-12-0381.

13. Nowotschin, S., Setty, M., Kuo, Y.-Y., Liu, V., Garg, V., Sharma, R., Simon, C.S., Saiz, N., Gardner, R., Boutet, S.C., et al. (2019). The emergent landscape of the mouse gut endoderm at single-cell resolution. Nature 569, 361–367. 10.1038/s41586-019-1127-1.

14. Kwon, G.S., Viotti, M., and Hadjantonakis, A.-K. (2008). The endoderm of the mouse embryo arises by dynamic widespread intercalation of embryonic and extraembryonic lineages. Dev. Cell 15, 509–520. 10.1016/j.devcel.2008.07.017.

15. Viotti, M., Nowotschin, S., and Hadjantonakis, A.-K. (2014). SOX17 links gut endoderm morphogenesis and germ layer segregation. Nat. Cell Biol. 16, 1146– 1156. 10.1038/ncb3070.

16. Thowfeequ, S., Fiorentino, J., Hu, D., Solovey, M., Ruane, S., Whitehead, M., Vanhaesebroeck, B., Scialdone, A., and Srinivas, S. (2021). Characterisation of the transcriptional dynamics underpinning the function, fate, and migration of the mouse Anterior Visceral Endoderm. bioRxiv, 2021.06.25.449902. 10.1101/2021.06.25.449902.

17. Mesnard, D., Filipe, M., Belo, J.A., and Zernicka-Goetz, M. (2004). The anterior-posterior axis emerges respecting the morphology of the mouse embryo that changes and aligns with the uterus before gastrulation. Curr. Biol. 14, 184–196. 10.1016/j.cub.2004.01.026.

18. Dimitrov, D., Türei, D., Garrido-Rodriguez, M., Burmedi, P.L., Nagai, J.S., Boys, C., Ramirez Flores, R.O., Kim, H., Szalai, B., Costa, I.G., et al. (2022). Comparison of methods and resources for cell-cell communication inference from single-cell RNA-Seq data. Nat. Commun. 13, 3224. 10.1038/s41467-022-30755-0.

19. Scheibner, K., Schirge, S., Burtscher, I., Büttner, M., Sterr, M., Yang, D., Böttcher, A., Ansarullah, Irmler, M., Beckers, J., et al. (2021). Epithelial cell plasticity drives endoderm formation during gastrulation. Nat. Cell Biol. 23, 692–703. 10.1038/s41556-021-00694-x.

20. Zhang, S., Chen, T., Chen, N., Gao, D., Shi, B., Kong, S., West, R.C., Yuan, Y., Zhi, M., Wei, Q., et al. (2019). Implantation initiation of self-assembled embryo-like structures generated using three types of mouse blastocyst-derived stem cells. Nat. Commun. 10, 496. 10.1038/s41467-019-08378-9.

21. Langkabel, J., Horne, A., Bonaguro, L., Holsten, L., Hesse, T., Knaus, A., Riedel, Y., Becker, M., Händler, K., Elmzzahi, T., et al. (2021). Induction of Rosette-to-Lumen stage embryoids using reprogramming paradigms in ESCs. Nat. Commun. 12, 7322. 10.1038/s41467-021-27586-w.

22. Vrij, E.J., Scholte op Reimer, Y.S., Fuentes, L.R., Guerreiro, I.M., Holzmann, V., Aldeguer, J.F., Sestini, G., Koo, B.-K., Kind, J., van Blitterswijk, C.A., et al. (2022). A pendulum of induction between the epiblast and extra-embryonic endoderm supports post-implantation progression. Development 149. 10.1242/dev.192310.

23. Bedzhov, I., and Zernicka-Goetz, M. (2014). Self-organizing properties of mouse pluripotent cells initiate morphogenesis upon implantation. Cell 156, 1032–1044. 10.1016/j.cell.2014.01.023.

24. Semrau, S., Goldmann, J.E., Soumillon, M., Mikkelsen, T.S., Jaenisch, R., and van Oudenaarden, A. (2017). Dynamics of lineage commitment revealed by single-cell transcriptomics of differentiating embryonic stem cells. Nat. Commun. 8, 1096. 10.1038/s41467-017-01076-4.

25. Lu, C.C., and Robertson, E.J. (2004). Multiple roles for Nodal in the epiblast of the mouse embryo in the establishment of anterior-posterior patterning. Dev. Biol. 273, 149–159. 10.1016/j.ydbio.2004.06.004.

26. Chazaud, C., and Rossant, J. (2006). Disruption of early proximodistal patterning and AVE formation in Apc mutants. Development 133, 3379–3387. 10.1242/dev.02523.

27. ten Berge, D., Kurek, D., Blauwkamp, T., Koole, W., Maas, A., Eroglu, E., Siu, R.K., and Nusse, R. (2011). Embryonic stem cells require Wnt proteins to prevent differentiation to epiblast stem cells. Nat. Cell Biol. 13, 1070–1075. 10.1038/ncb2314.

28. Neagu, A., van Genderen, E., Escudero, I., Verwegen, L., Kurek, D., Lehmann, J., Stel, J., Dirks, R.A.M., van Mierlo, G., Maas, A., et al. (2020). In vitro capture and characterization of embryonic rosette-stage pluripotency between naive and primed states. Nat. Cell Biol. 22, 534–545. 10.1038/s41556-020-0508-x.

29. Fan, R., Kim, Y.S., Wu, J., Chen, R., Zeuschner, D., Mildner, K., Adachi, K., Wu, G., Galatidou, S., Li, J., et al. (2020). Wnt/Beta-catenin/Esrrb signalling controls the tissue-scale reorganization and maintenance of the pluripotent lineage during murine embryonic diapause. Nat. Commun. 11, 5499. 10.1038/s41467-020-19353-0.

30. Sick, S., Reinker, S., Timmer, J., and Schlake, T. (2006). WNT and DKK determine hair follicle spacing through a reaction-diffusion mechanism. Science 314, 1447–1450. 10.1126/science.1130088.

31. T, S., JP, C., S, W., H, T.-K.V., R, B., SY, L., B, F., F, J., and JC, R. (2017). Antagonistic Self-Organizing Patterning Systems Control Maintenance and Regeneration of the Anteroposterior Axis in Planarians. Dev. Cell 40. 10.1016/J.DEVCEL.2016.12.024.

32. Hooper, M., Hardy, K., Handyside, A., Hunter, S., and Monk, M. (1987). HPRT-deficient (Lesch-Nyhan) mouse embryos derived from germline colonization by cultured cells. Nature 326, 292–295. 10.1038/326292a0.

33. Ying, Q.-L., Wray, J., Nichols, J., Batlle-Morera, L., Doble, B., Woodgett, J., Cohen, P., and Smith, A. (2008). The ground state of embryonic stem cell self-renewal. Nature 453, 519–523. 10.1038/nature06968.

34. Morgani, S.M., Saiz, N., Garg, V., Raina, D., Simon, C.S., Kang, M., Arias, A.M., Nichols, J., Schröter, C., and Hadjantonakis, A.-K. (2018). A Sprouty4 reporter to monitor FGF/ERK signaling activity in ESCs and mice. Dev. Biol. 441, 104–126. 10.1016/j.ydbio.2018.06.017.

35. Wang, W., Lin, C., Lu, D., Ning, Z., Cox, T., Melvin, D., Wang, X., Bradley, A., and Liu, P. (2008). Chromosomal transposition of *PiggyBac* in mouse embryonic stem cells. Proc. Natl. Acad. Sci. 105, 9290–9295. 10.1073/pnas.0801017105.

36. Ran, F.A., Hsu, P.D., Wright, J., Agarwala, V., Scott, D.A., and Zhang, F. (2013). Genome engineering using the CRISPR-Cas9 system. Nat. Protoc. 8, 2281. 10.1038/NPROT.2013.143.

37. Kim, Y.S., Fan, R., Kremer, L., Kuempel-Rink, N., Mildner, K., Zeuschner, D., Hekking, L., Stehling, M., and Bedzhov, I. (2021). Deciphering epiblast lumenogenesis reveals proamniotic cavity control of embryo growth and patterning. Sci. Adv. 7. 10.1126/sciadv.abe1640.

38. Schröter, C., Rué, P., Mackenzie, J.P., and Arias, A.M. (2015). FGF/MAPK signaling sets the switching threshold of a bistable circuit controlling cell fate decisions in ES cells. Development 142, 4205–4216. 10.1242/dev.127530.

39. Choi, H.M.T., Schwarzkopf, M., Fornace, M.E., Acharya, A., Artavanis, G., Stegmaier, J., Cunha, A., and Pierce, N.A. (2018). Third-generation in situ hybridization chain reaction: multiplexed, quantitative, sensitive, versatile, robust. Development 145. 10.1242/DEV.165753.

40. Edelstein, A., Amodaj, N., Hoover, K., Vale, R., and Stuurman, N. (2010). Computer Control of Microscopes Using µManager. Curr. Protoc. Mol. Biol. 92, 14.20.1-14.20.17. 10.1002/0471142727.mb1420s92.

41. Preibisch, S., Saalfeld, S., and Tomancak, P. (2009). Globally optimal stitching of tiled 3D microscopic image acquisitions. Bioinformatics 25, 1463–1465. 10.1093/bioinformatics/btp184.

42. Hao, Y., Hao, S., Andersen-Nissen, E., Mauck, W.M., Zheng, S., Butler, A., Lee, M.J., Wilk, A.J., Darby, C., Zager, M., et al. (2021). Integrated analysis of multimodal single-cell data. Cell 184, 3573–3587.e29. 10.1016/j.cell.2021.04.048.

43. Wolf, F.A., Angerer, P., and Theis, F.J. (2018). SCANPY: large-scale single-cell gene expression data analysis. Genome Biol. 19, 15. 10.1186/s13059-017-1382-0.

